# Reducing chlorophyll level in seed filling stages results in higher seed nitrogen without impacting canopy carbon assimilation

**DOI:** 10.1101/2022.07.01.498509

**Authors:** Young B. Cho, Ryan A. Boyd, Yudong Ren, Moon-Sub Lee, Sarah I. Jones, Ursula M. Ruiz-Vera, Justin McGrath, Michael D. Masters, Donald R. Ort

## Abstract

Chlorophyll is the major light absorbing pigment for plant photosynthesis. While evolution has selected for higher chlorophyll content in leaves, previous work suggests that domesticated crops grown in modern agricultural environments overinvest in chlorophyll production thereby lowering light use and nitrogen use efficiency. To investigate the potential benefits of reducing chlorophyll level, we created ethanol inducible RNAi tobacco mutants that suppress Mg-chelatase subunit I (*CHLI*) with small RNA (sRNA) within 3 hours of induction and reduce chlorophyll within 5 days in field conditions. We initiated chlorophyll reduction later in plant development to avoid the highly sensitive seedling stage, and to allow young plants to have full green leaves to maximize light interception before canopy formation. This study demonstrated that >60% reduction of leaf chlorophyll could be tolerated without penalty on above-ground biomass or canopy photosynthesis in field conditions. Leaf chlorophyll reduction during seed filling stages increased tobacco seed nitrogen concentration as much as 17%, while biomass and seed yields were maintained. These results indicate that time-specific reduction of chlorophyll could be a niche strategy that decouples the inverse relationship between yield and seed nitrogen by utilizing saved nitrogen from the reduction of chlorophyll while maintaining full carbon assimilation capacity.

## INTRODUCTION

Closing the gap between yield potential and realized current yield of crops while maintaining nutritional quality is necessary to meet future global food demand (Zhu et al., 2010; Long and Ort, 2010; Ray et al., 2013). A significant component of the yield gap is the lower theoretical efficiency of photosynthesis (Ainsworth and Long, 2021; Ort et al., 2015). The efficiency of photosynthesis often decreases when the amount of absorbed photosynthetically active radiation increases (Sinclair and Muchow, 1999; Slattery et al., 2013; Slattery and Ort, 2015), suggesting that for agricultural purposes crop plants invest too many resources in light capture while underinvesting in light utilization. A major evolutionary benefit of overinvestment in light capture was that shading potential competitors conferred a selective advantage (Zhang et al., 1999). Even when photosynthesis is light-saturated and thus cannot utilize additional light, intercepting more light prevents a potential competitor from receiving and benefitting from the light. However, this investment strategy is suboptimal for crops in a monoculture environment (Loomis, 1993; Denison et al., 2003) where the goal is to maximize net primary productivity for the field. In agricultural canopies, radiation penetration, and therefore radiation use efficiency, is decreased by dense foliage at the top of the canopy, which absorbs most of the incident photosynthetically active radiation (Long, 1993; Long et al., 2006). The rate of photosynthesis reaches saturation (*Asat*) at a light intensity well below that of full sunlight. At moderate to high light intensities, the rate at which sunlight is absorbed by sun-exposed leaves vastly exceeds the amount needed to reach *Asat*, and the excess absorbed photons are dissipated by photoprotective mechanisms and thereby wasted.

Modeling studies have proposed that plants produce excess chlorophyll and thus restrained production of chlorophyll could benefit nitrogen use efficiency without compromising total canopy photosynthesis. Using the sunlit-shaded model, Ort et al. (2011) proposed that a 50% reduction in chlorophyll would improve light distribution and increase canopy photosynthesis; however, further decreases in chlorophyll would be disadvantageous. Studies using advanced multi-layer canopy models have predicted that leaf chlorophyll could be decreased by 50% without penalty to canopy photosynthesis, while additionally bringing about a potential 9% savings of leaf nitrogen (Walker et al., 2018). A separate modeling study concluded that a 60% reduction in chlorophyll could increase nitrogen use efficiency and increase canopy photosynthesis if the saved nitrogen could be reinvested to increase photosynthetic capacity in areas of the canopy where light intensity was increased (Song et al., 2017). In addition, experimental evidence supports the premise that lowering leaf chlorophyll could be beneficial. Field-grown low chlorophyll rice and soybean show similar or greater photosystem II efficiency, Rubisco carboxylation rates, and nitrogen-use efficiency when compared to dark green wild-types (Li et al., 2013; Gu et al., 2017; Sakowska et al., 2018). Low chlorophyll rice and tobacco mutants showed higher yields than dark green wild-types at higher planting densities (Gu et al., 2017; Kirst et al., 2017). However, one predicted disadvantage of reduced leaf chlorophyll is that low chlorophyll content early in the season before canopy closure will result in reduced light interception efficiency because less light is absorbed by young leaves while more light is transmitted to the soil. Canopy modeling predicts increased canopy photosynthesis after canopy closure and decreased photosynthesis before the closure (Long et al., 2006; Song et al., 2017). The benefits of lowering chlorophyll are expected to depend on both the timing and extent of chlorophyll reduction, which can be realized by time-specific gene regulation through inducible promoters. The ethanol inducible gene expression system (Felenbok et al., 1988) found in fungi has been adopted in plant science (Caddick et al., 1998; Salter et al., 1998) and has previously been used to reduce chlorophyll synthesis in tobacco (Chen et al., 2003) thus, it is a practical tool for realizing time-specific chlorophyll regulation.

Reducing chlorophyll production could free nitrogen resources for other uses by the plant, possibly increasing seed nitrogen without compromising carbon assimilation. In cereals, the nitrogen content in seeds reflects the seed protein content and thus the seed nutritional quality (Good et al., 2004). Studies about the genetic basis for seed composition have revealed that most cereals (Simmonds, 1995), including maize (Feil et al., 1990), wheat (Canevara et al., 1994), and oilseed rape (Brennan et al., 2000), have an inverse relationship between seed yield and protein concentration. This occurs because of the dilution of proteins by carbohydrates (Acreche and Slafer, 2009) and the competition between carbon and nitrogen metabolisms for energy (Munier-Jolain and Salon, 2005). Plants can use leaves for either carbon assimilation or nitrogen remobilization, which are mutually exclusive processes (Havé et al., 2017). Leaves capture solar energy for photosynthetic carbon assimilation and export carbon in the form of sugars to the seeds. Reduced photosynthetic activity during senescence decreases carbon assimilation and sugar export to seeds. On the other hand, nitrogen is transferred from leaves to seeds during senescence, as protein is broken down (Masclaux-Daubresse et al., 2008). Delaying leaf senescence increases seed yield due to the maintenance of carbon assimilation, while the maintenance of the photosynthetic apparatus delays nitrogen remobilization and decreases seed N content. For cereals and crops that are cultivated for their seed protein content, this constitutes a dilemma that opposes yield performance against seed nitrogen (Good et al., 2004; Uauy et al., 2006; Oury and Godin, 2007; Gregersen et al., 2013; Distelfeld et al., 2014). Perhaps, saved nitrogen from decreased chlorophyll production could increase seed nitrogen without penalty on yield because more free nitrogen is available in the seed filling stages without compromising canopy photosynthesis, as models have predicted (Ort et al., 2011; Walker et al., 2018; Song et al., 2017).

The main objective of this study was to investigate the impact of reducing chlorophyll in a time-specific manner, particularly in later plant developmental stages after canopy closure and at the start of seed filling using a combination of field and greenhouse work. We hypothesized that lowering chlorophyll at a later developmental stage would not decrease canopy photosynthesis, biomass, and seed yield. We predict the reduction of leaf N sequestration will increase seed nitrogen without a reduction in net carbon assimilation. To test these hypotheses we used transformed tobacco (cv. Petite Havana), which generated small RNA (sRNA) that down-regulated Mg-chelatase subunit I (*CHLI*) expression via an ethanol inducible promoter, leading to the reduction of chlorophyll only when ethanol was applied. We initiated chlorophyll reduction later in the plant’s growth stage at the start of seed filling which allowed young plants to have full green leaves to maximize light interception before the canopy was formed, when losses to light inhomogeneity were small. This study demonstrated that up to 62% reduction of leaf chlorophyll after canopy closure was possible without penalty to above-ground biomass or on leaf and canopy photosynthesis under field conditions. Moreover, leaf chlorophyll reduction initiated at the start of seed filling resulted in as much as a 17% increase in seed nitrogen concentration with no reduction in biomass or seed yield.

## RESULTS

### Ethanol inducible RNAi mutant can suppress chlorophyll levels by down-regulating Mg-chelatase subunit I in a developmental stage specific manner

To decrease leaf chlorophyll we designed an ethanol inducible RNAi construct that targets the Mg-chelatase subunit I (*CHLI*) (Nitab4.5_0006237g0030). Mg-chelatase is an effective target to reduce chlorophyll synthesis (Figure 1A). Down-regulation of *CHLI* avoids the excessive accumulation of highly photoreactive and cytotoxic chlorophyll precursors that could occur if other enzymes of the pathway were targeted. There are naturally occurring as well as mutagenized low chlorophyll soybean mutants with genetic disruption in the Mg-chelatase enzyme (Campbell et al., 2015) that grow as strongly as the isogenic wild-type parent (Slattery et al., 2017). The 35S promoter constitutively expresses the *Alc* regulon, which binds to the *alcA* promoter in the presence of ethanol (Figure 1B). We inserted the ethanol inducible *CHLI* RNAi construct into *Nicotiana tabacum* (cv. Petit Havana) and selected mutant1 (mt1) and mutant2 (mt2) which responded to ethanol treatment among 22 transformation events. The T-DNA is located on scaffold 161: 578931 in mt1 and on scaffold 638:555,535 in mt2 (Figure 1C, Supplemental Figures 1 and 2).

**Figure 1.**
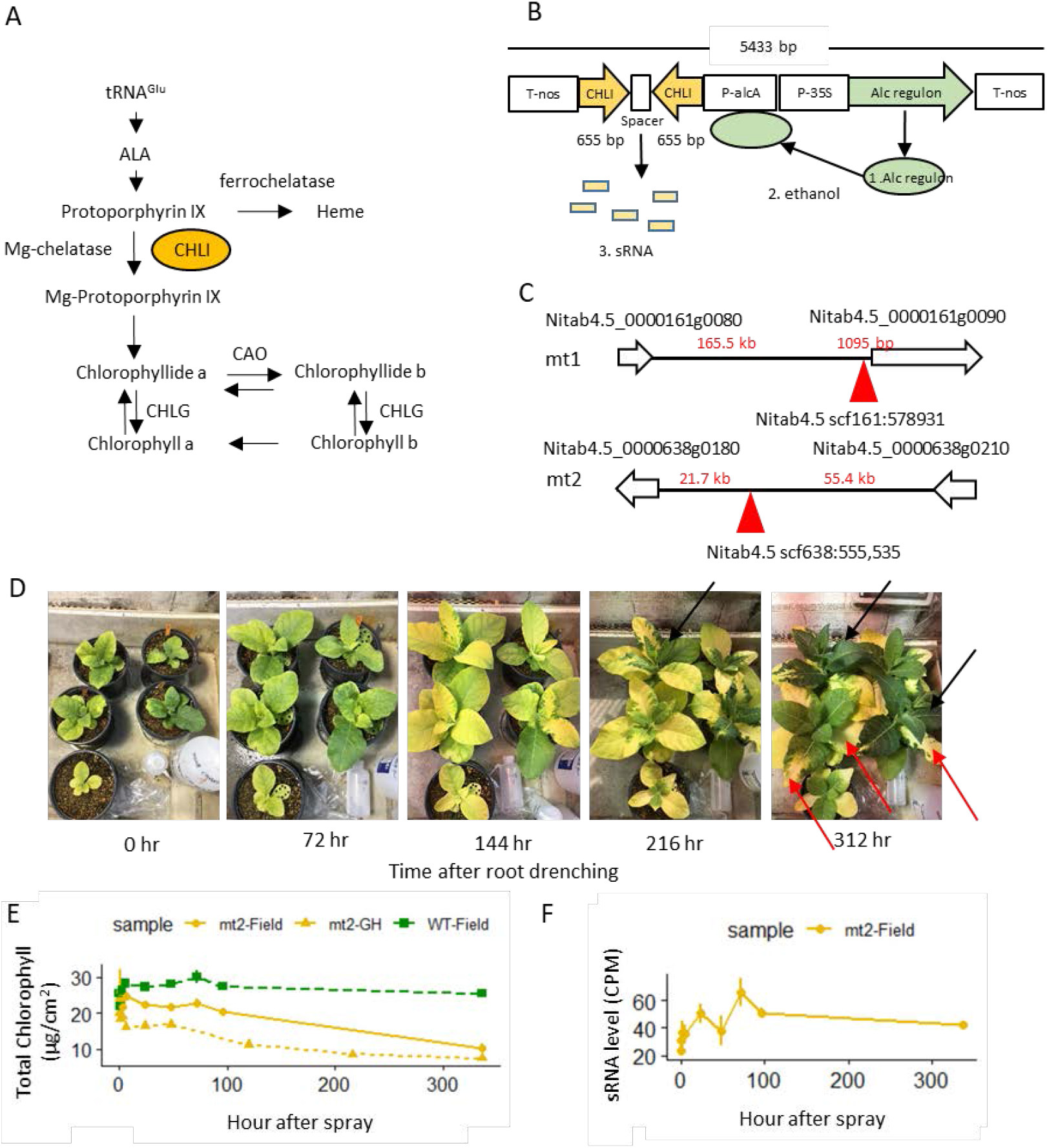
Developmental stage specific down-regulation of chlorophyll using ethanol inducible promoter. A. Biosynthesis pathway of chlorophyll. B. Design of the sRNA construct. C. Location of T-DNA in the tobacco genome. T-DNA (red triangle) is located at scaffold 161:578931 in mt1 and at scaffold 638:555,535 in mt2. D. Time specific regulation of chlorophyll. 100ml 1% ethanol applied to mt1 roots in the greenhouse. 216 hr after induction, newly developed leaves are fully dark green (black arrow) while affected leaves and partial leaf area (red arrow) did not recover from the low chlorophyll phenotype. E-F. Levels of total chlorophyll and sRNA expression after 2% ethanol spray to leaves of wild-type and mt2 in the greenhouse and 2019 Illinois field. Error bars represent standard error (n=4). E. The level of chlorophyll. F. The level of CHLI sRNA expression (CPM, counts per million).

We tested the response of the mutants to ethanol treatment by adding 100 ml of 1% (v/v) ethanol to the roots of mt1 in the greenhouse. Leaves began to exhibit a low chlorophyll phenotype 72 hours (3 days) after ethanol application (Figure 1D). Leaves that developed 216 hours (9 days) after the ethanol application were completely dark green (Figure 1D, black arrows), indicating that the plants reinitiated chlorophyll synthesis once the effects of the RNA interference abated. In some cases, the whole leaf or partial area of a leaf (Figure 1D, red arrows) did not recover from the low chlorophyll phenotype even though the plant reinitiated chlorophyll synthesis in new leaves (Figure 1D). We determined this root drenching method had two major issues that made it impractical for experimental trials. First, chlorophyll in existing leaves was so greatly reduced that the leaves turned completely white; canopy modeling suggested that greater than a 70% reduction in chlorophyll would not be beneficial for canopy photosynthesis (Ort et al., 2011). Second, root drenching would not be practical for large-scale field trials. To overcome these limitations, we tested canopy spray methods in a field setting. We sprayed 2% ethanol on WT, mt1, and mt2 leaves in the field every morning (between 7-9 AM) until leaves were soaked with the solution. The level of chlorophyll of mt2 began to decrease 120 hours (5 days) after the first ethanol spray (Figure 1E), while the wildtype chlorophyll was unaffected. The sRNA expression increased 3 hours after the first ethanol spray (Figure 1F), peaked at 72 hours (3 days) and remained at a high level throughout the daily treatment regimen. In summary, this spray methodology effectively reduced chlorophyll to a desirable level in the mutants after the canopy was closed, avoiding the disadvantages of lowering chlorophyll in early development stages.

### Up to 62% chlorophyll reduction in later development stages resulted in no change of above-ground biomass in the field tests

To understand the impact of reduced chlorophyll level on canopy photosynthesis and biomass, we tested the ethanol inducible RNAi mutants in three tobacco field trials; Illinois in 2019 (IL2019), Puerto Rico in 2019 (PR2019), and Illinois in 2020 (IL2020). In the Illinois 2019 field experiment (IL2019), we tested three genotypes: wild-type (WT), mutant1 (mt1), and mutant2 (mt2). We had three treatments per genotype--spray once a day (-1), spray twice a day (-2), or no treatment (−0)--to evaluate the effect of ethanol spray on chlorophyll level of canopy and above-ground biomass. The plants were sprayed with 2% (v/v) ethanol from canopy closure (48 DAP, days after planting) until one week before harvest (74 DAP). The low chlorophyll phenotype was easily discernible with the naked eye (Figure 2A). We measured the level of chlorophyll of the canopy during the entire field season. Chlorophyll level in mt2-2 (sprayed twice a day) plot began to decrease after 5 days of ethanol spraying and remained 30-50% lower than the wild-type plot during the remainder of the season (Figure 2C, Supplemental Table 1). Despite the reduction in chlorophyll, we observed no change in aboveground biomass between mt2 treatments (Fig 2B). Unlike the response that we observed in the greenhouse, mt1 did not respond to ethanol induction in the field, such that mt1-0, mt1-1, and mt1-2 showed no significant difference in chlorophyll level. Dried above-ground biomass showed no significant difference between genotypes and treatments (Figure 2B). In summary, the ethanol spray did not affect the above-ground biomass or chlorophyll level of mt1 and WT genotypes. In contrast, the ethanol spray reduced the chlorophyll levels of mt2 (mt2-1 and mt2-2) by up to 50% while no change was observed in the above-ground biomass.

**Figure 2.**
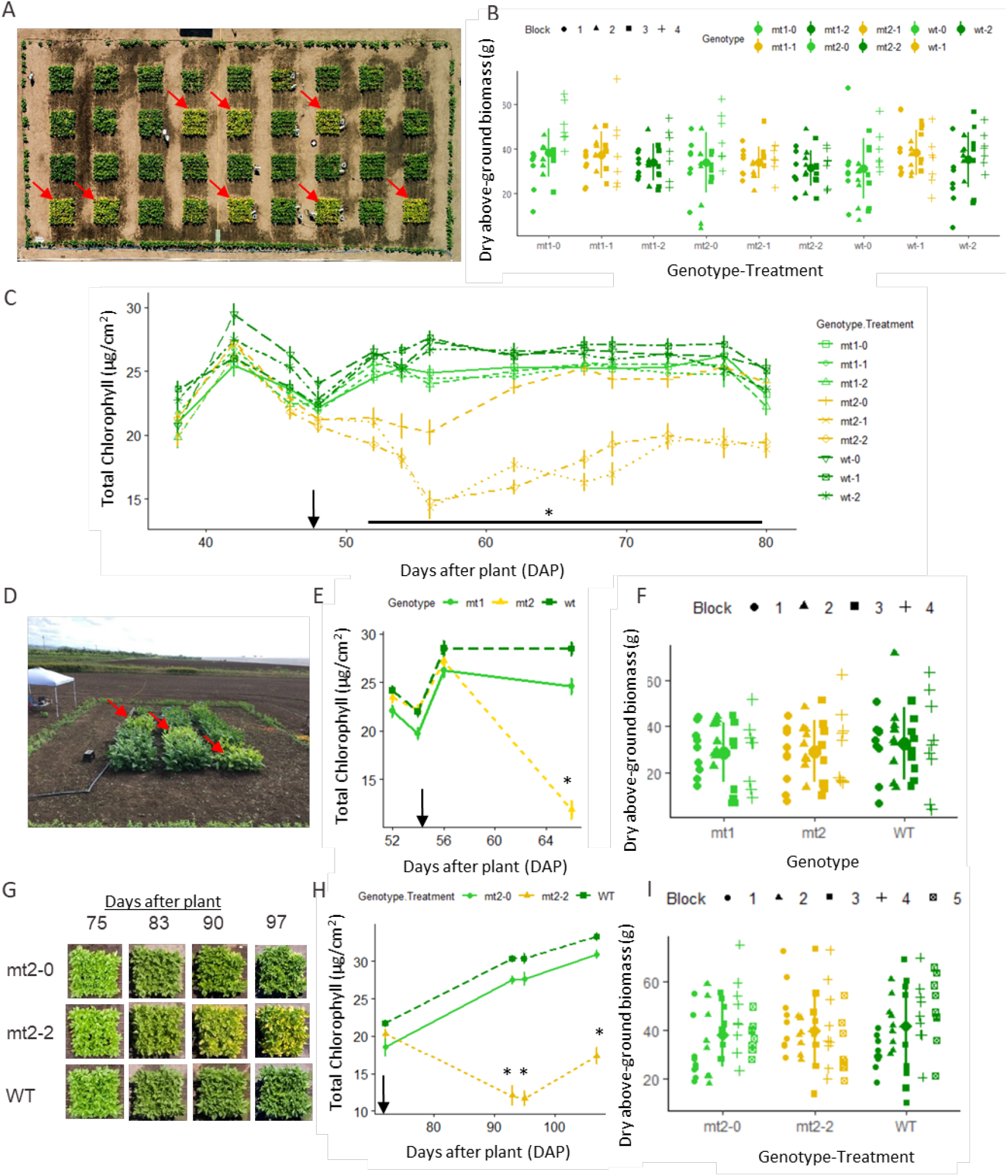
Up to 62% chlorophyll reduction after canopy closure results in no change on above-ground biomass in the field tests. A-C. 2019 Illinois field experiment (n=4 blocks). Three genotypes--wild-type (WT), mutant1 (mt1), and mutant2 (mt2)--were treated with three treatments: no spray (−0), 2% ethanol spray once daily (-1), twice daily (-2). D-F. 2019 Puerto Rico field experiment (n=4 blocks). Three genotypes--WT, mt1, and mt2--were treated with 2% ethanol spray twice a day. G-I. 2020 Illinois field experiment (n=5 blocks) has two genotypes, WT and mt2, which were treated with 2% ethanol spray twice a day, in addition to mt2 without ethanol treatment (mt2-0). A, D and G. Aerial view of the field trial shows the reduction of chlorophyll in the treated mt2 plots (red arrow). B, F and I. Dry above ground biomass.Each dot represents a subsample within a block. Error bars represent standard deviations of blocks, not subsamples. C, E and H. Chlorophyll level of sun leaves during field experiment. Error bars represent standard errors. Black arrow indicates when spraying started. The statistical analysis was done using linear mixed model and type III ANOVA (n= number of block). * indicates significant differences between treated mt2 and WT within DAP when present (p<0.05).

The IL2019 trial revealed no effect of sprayed ethanol on the above-ground biomass of full green genotypes, so we focused on the comparison of chlorophyll reduction and above-ground biomass in successive field trials. To assess this effect, three genotypes (WT, mt1, and mt2) were tested with one spray treatment, twice daily with 2% (v/v) ethanol in Puerto Rico in 2019 (PR2019). A 58% decrease in canopy chlorophyll level was observed in the mt2 genotype after treatment (Figures 2D and E, Supplemental Table 1), while the above-ground biomass was not significantly different (Figure 2F). In the IL2020 field trial (IL2020), only WT and mt2 genotypes were used as mt1 plants did not respond to the sprayed ethanol treatment and had similar chlorophyll content as WT plants in previous field trials. In an effort to further elucidate the relationship between the Mg chelatase mutation and chlorophyll reduction we added an mt2 treatment without sprayed ethanol induction to this field trial. There was no difference in above-ground biomass between any treatment (WT-2, mt2-2, mt2-0, Figure 2I), while up to a 62% chlorophyll reduction was observed in mt2-2 (Figures 2G and H, Supplemental Table 1).

Cumulatively the evidence from these three field trials demonstrates that a twice daily canopy spraying of ethanol effectively reduced chlorophyll content in mt2 tobacco by up to 62% with the magnitude of the response varying between year and location. Perhaps more importantly these trials have demonstrated with repeatability that large reductions in chlorophyll after canopy closure had no effect on the above-ground biomass of tobacco. [We did not collect seeds from any of these field trials due to APHIS (Animal and Plant Health Inspection Service) regulations regarding seed dispersal.]

### Reducing chlorophyll content of leaves did not significantly change canopy carbon assimilation rate

To assess the impact of reduced chlorophyll level on leaf and canopy photosynthesis, the absorbed light and CO_2_ response of net CO_2_ assimilation rate (*A*_net_) were measured on sun exposed leaves in three field seasons [IL2019, PR2019, and IL2020 (Sun)]. Additionally, a shade leaf (4 nodes below sun leaf) was also measured for a single field season [IL2020 (Shade)] to assess gas exchange in lower canopy leaves. Despite the significant reduction in chlorophyll levels observed in all mt2 field trials, we observed minimal differences between low chlorophyll leaves and full green leaves for all photosynthesis parameters in the light response curves (*A*/*Q*_abs_ curves; Figures 3A-D). The respiration rate (*R*), the light saturated rate of net CO_2_ assimilation (*A*_satQ_), and the empirical curvature factor (*θ*) were not different between low chlorophyll and full green leaves for any of the *A*/*Q*_abs_ curve data sets (Table 1). Reductions in the maximum quantum yield of CO_2_ assimilation (*Φ_CO2_*) were observed in IL2019 and PR2019 experiments for the low chlorophyll, but not in IL2020 (Table 1). For the IL2020 field season, the maximal photochemical yield of photosystem II obtained at low light (*Φ*_PSII(LL)_) was lower in leaves with low chlorophyll content, but this was not consistently observed in the other field trials (Supplemental Table 2). The response of *A*_net_ to internal CO_2_ concentration (*C*_i_; *A*/*C*_i_ curves; Figures 3E-H) showed inconsistent differences in photosynthetic parameters, while the empirical curvature factors for the *A*/*C*_i_ curves (*ω*) were consistently lower in low chlorophyll leaves (Supplemental Table 2). Total net carbon assimilation of the canopy (*A*′) was calculated by using a simple three-leaf level concept (Supplemental Figure 4) simulating an LAI (leaf area index) of 3 similar to measurements from the field trial (Supplemental Figure 5). No difference in canopy photosynthesis between low chlorophyll and dark green canopies (Table 1) were calculated for all field trials. We used IL2020 gas exchange data measured from the shade leaf (4 nodes below sun leaf) for the second and third levels in the canopy calculation; IL2020 sun leaf gas exchange data were used for the uppermost leaf layer in the calculation. *A*′ of IL2020 shade data were not significantly different between low chlorophyll and dark green canopies (Table 1). Overall, we did not observe significant differences in canopy photosynthesis between low chlorophyll and dark green controls in any of the three field experiments.

**Figure 3.**
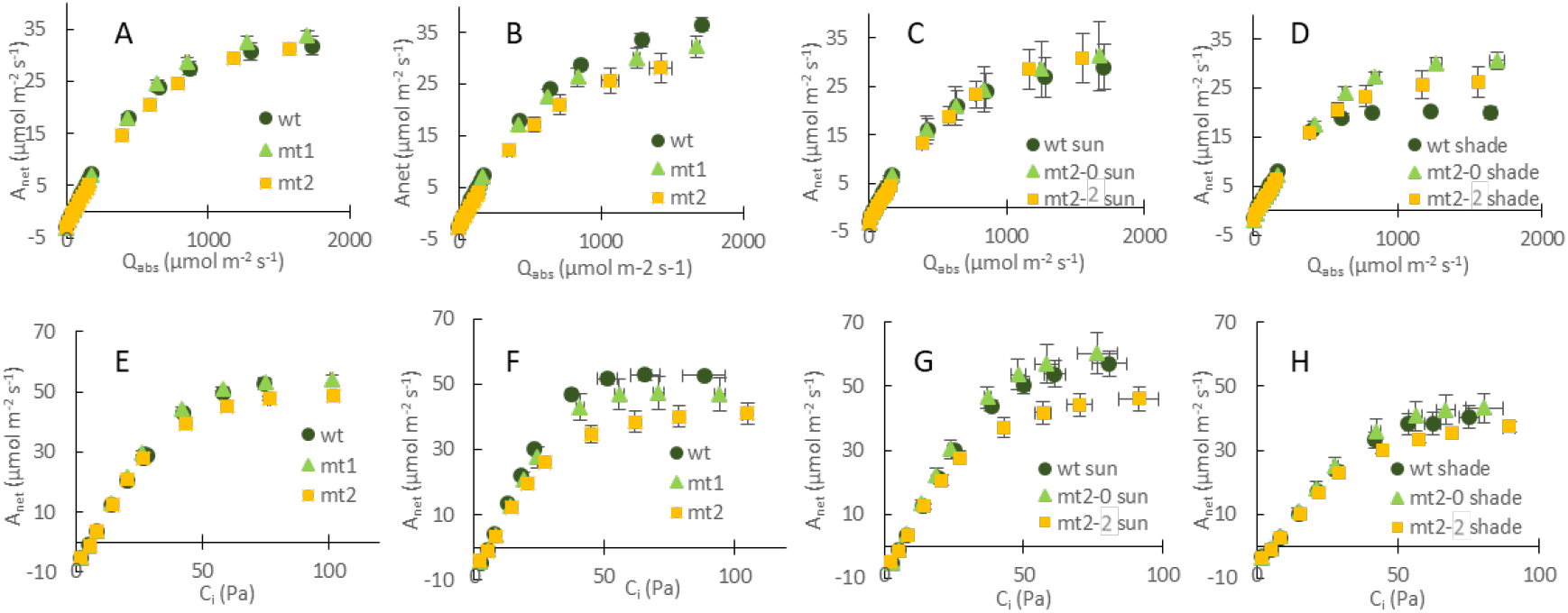
Photosynthetic capacity of low and dark green controls from three field trials. Panels A through D show the light response curves. Panels E through H show the CO_2_ response curves. A and E, IL2019 (n=4); B and F, PR2019 (n=3); C and G, IL2020 sun leaf (n=4); D and H, IL2020 shade leaf (n=3). IL2019 data (n=4) and PR2019 (n=3) compared genotypes within the treatment, 2% ethanol spray twice a day. IL2020 (n=4) has two genotypes, WT and mt2, which were treated with 2% ethanol spray twice a day, in addition to mt2 without ethanol treatment (mt2-0). Shapes indicate the mean of three to four field plots with ± standard error.

**Table 1.**
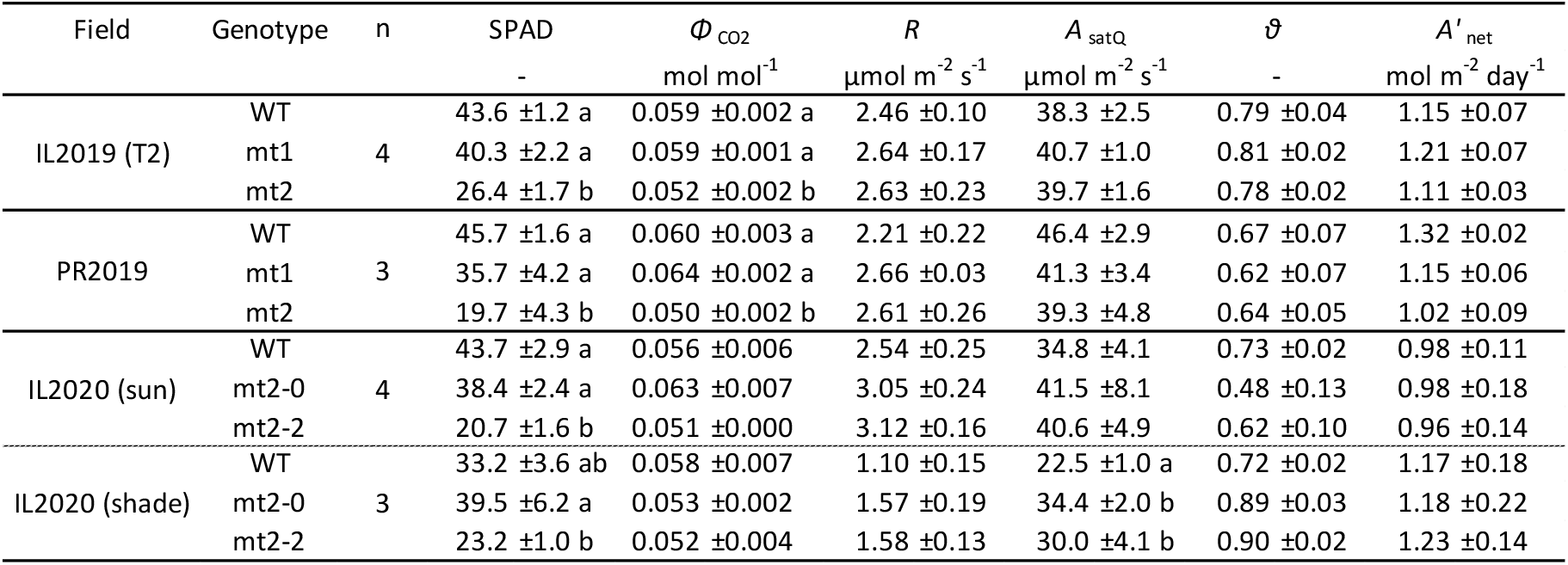
Low and high chlorophyll leaf pigment and photosynthetic parameters from three field trials. The results were determined from *A/Q_abs_* curves (Figure 3) after fitting the data to a non-rectangular hyperbola. The maximum quantum yield of CO_2_ assimilation (*Φ_CO2_*), respiration rate (*R*), the light saturated rate of net CO_2_ assimilation (*A_sat_*), and the empirical curvature factor (*θ*) were calculated. The statistical analysis was done using linear mixed model and type III ANOVA with post-hoc Tukey tests. IL2019 data (n=4) compared genotypes within the treatment (T2). PR2019 (n=3). IL2020 (n=4). Means ± standard errors are shown. Different letters represent significant differences (p < 0.05).

| Field | Genotype | n | SPAD | $\Phi_{CO_2}$<br>mol mol<sup>-1</sup> | $R$<br>μmol m<sup>-2</sup> s<sup>-1</sup> | $A_{satQ}$<br>μmol m<sup>-2</sup> s<sup>-1</sup> | $\theta$<br>- | $A'_{net}$<br>mol m<sup>-2</sup> day<sup>-1</sup> |
| --- | --- | --- | --- | --- | --- | --- | --- | --- |
| IL2019 (T2) | WT | 4 | 43.6 ±1.2 a | 0.059 ±0.002 a | 2.46 ±0.10 | 38.3 ±2.5 | 0.79 ±0.04 | 1.15 ±0.07 |
|  | mt1 |  | 40.3 ±2.2 a | 0.059 ±0.001 a | 2.64 ±0.17 | 40.7 ±1.0 | 0.81 ±0.02 | 1.21 ±0.07 |
|  | mt2 |  | 26.4 ±1.7 b | 0.052 ±0.002 b | 2.63 ±0.23 | 39.7 ±1.6 | 0.78 ±0.02 | 1.11 ±0.03 |
| PR2019 | WT | 3 | 45.7 ±1.6 a | 0.060 ±0.003 a | 2.21 ±0.22 | 46.4 ±2.9 | 0.67 ±0.07 | 1.32 ±0.02 |
|  | mt1 |  | 35.7 ±4.2 a | 0.064 ±0.002 a | 2.66 ±0.03 | 41.3 ±3.4 | 0.62 ±0.07 | 1.15 ±0.06 |
|  | mt2 |  | 19.7 ±4.3 b | 0.050 ±0.002 b | 2.61 ±0.26 | 39.3 ±4.8 | 0.64 ±0.05 | 1.02 ±0.09 |
| IL2020 (sun) | WT | 4 | 43.7 ±2.9 a | 0.056 ±0.006 | 2.54 ±0.25 | 34.8 ±4.1 | 0.73 ±0.02 | 0.98 ±0.11 |
|  | mt2-0 |  | 38.4 ±2.4 a | 0.063 ±0.007 | 3.05 ±0.24 | 41.5 ±8.1 | 0.48 ±0.13 | 0.98 ±0.18 |
|  | mt2-2 |  | 20.7 ±1.6 b | 0.051 ±0.000 | 3.12 ±0.16 | 40.6 ±4.9 | 0.62 ±0.10 | 0.96 ±0.14 |
| IL2020 (shade) | WT | 3 | 33.2 ±3.6 ab | 0.058 ±0.007 | 1.10 ±0.15 | 22.5 ±1.0 a | 0.72 ±0.02 | 1.17 ±0.18 |
|  | mt2-0 |  | 39.5 ±6.2 a | 0.053 ±0.002 | 1.57 ±0.19 | 34.4 ±2.0 b | 0.89 ±0.03 | 1.18 ±0.22 |
|  | mt2-2 |  | 23.2 ±1.0 b | 0.052 ±0.004 | 1.58 ±0.13 | 30.0 ±4.1 b | 0.90 ±0.02 | 1.23 ±0.14 |

### Reducing chlorophyll level of leaves in the seed filling stages resulted in higher nitrogen concentration in the seed without penalty to biomass or seed yield

In the greenhouse, we tested the hypothesis that N saved from chlorophyll production could be used for seed protein by subjecting the inducible RNAi tobacco mutant (mt2) to variable soil N: we chose four levels of insufficient nitrogen and one with an adequate supply. The 2% (v/v) ethanol spray started 60 DAP, when flowering began (followed shortly by seed filling). The levels of chlorophyll at 84 DAP were significantly lower in the ethanol sprayed mutants than the WT in all treatments (Figure 4A). Chlorophyll reduction in mutants was between 60% to 68% less in the nitrogen deficient conditions, and 54% less in the adequate nitrogen condition. Plants were harvested at 101 DAP, and plant material was collected to measure dry biomass, and seeds were collected for weight and tissue composition. There was no difference in above-ground biomass or seed weight between WT and mutants for all treatments (Figures 4B and C). However, the nitrogen concentration was higher in low chlorophyll mutants under the 0.8 g urea (deficient) and Osmocote (adequate) conditions by 17% and 4% respectively (Fig 4D). Seed carbon concentration remained the same in all treatments (Figure 4E). However, due to the increase in nitrogen, the carbon to nitrogen ratio (C/N) of the seeds was reduced by 8% in nitrogen deficient and 4% in nitrogen sufficient conditions (Figure 4F).

**Figure 4.**
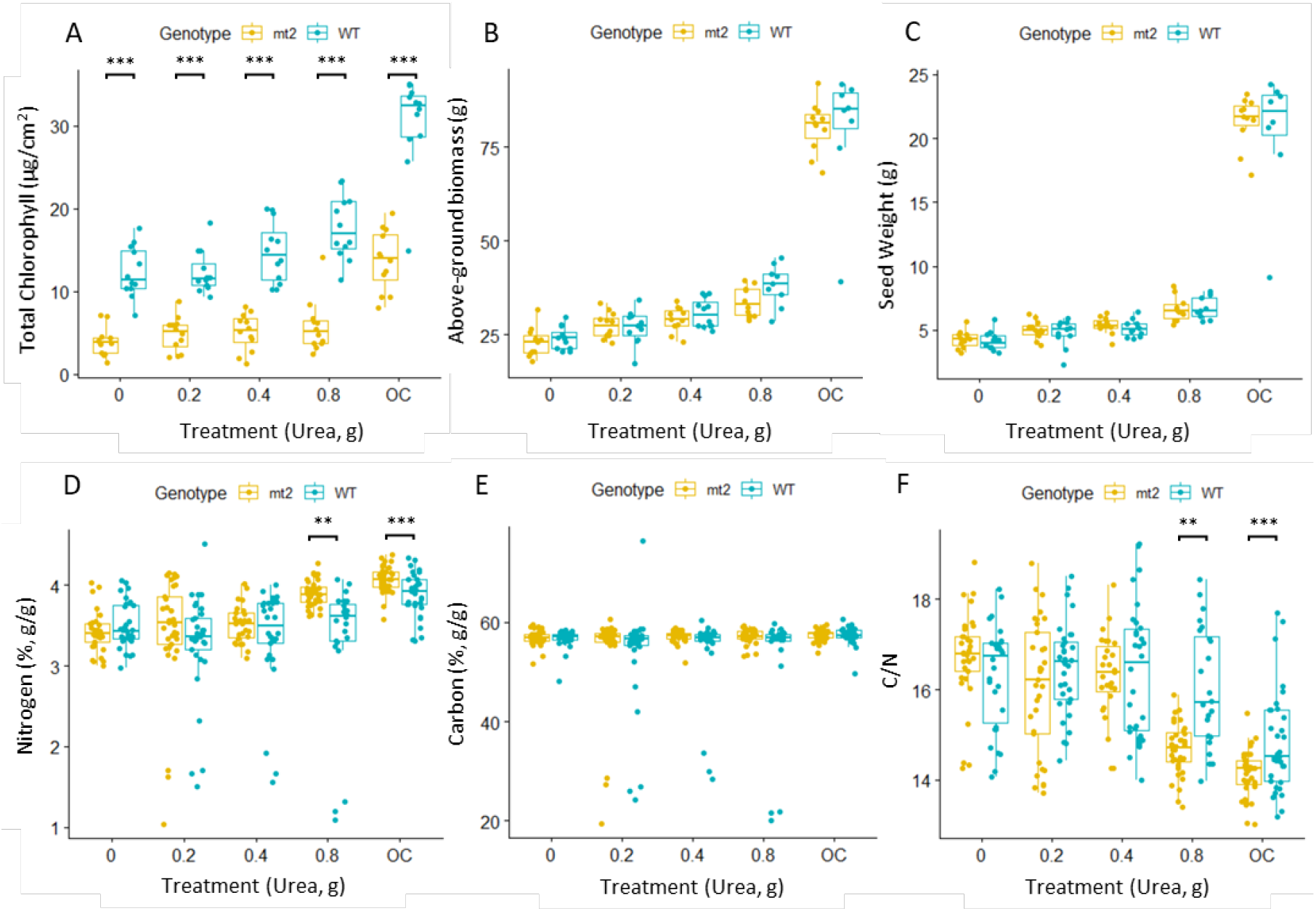
Increased nitrogen concentration and low C/N in seeds of low chlorophyll mutants in the greenhouse. Wild type (WT) and mutant2 (mt2) tobacco treated with different amounts of fertilizer in the greenhouse. X-axis shows the amount of urea added in a pot. OC stands for one teaspoon of Osmocote. A. Total chlorophyll level 24 days after treatment (84 DAP, n=12). B. Above ground biomass (n=12). C. Seed weight (n=12). D. Nitrogen concentration of seed (n=36). E. Carbon concentration of seed (n=36). F. C/N ratio of seed (n=36). The box plots show the median (central line), the lower and upper quartiles (box), and the minimum and maximum values (whiskers). Stars indicate significant difference within treatment when present (*, p<0.05; **, p<0.01; ***, p<0.001). The statistical analysis was done using the Wilcoxon rank test.

Under field conditions using conventional fertilization practices, we evaluated the impact of reducing the chlorophyll level of the leaves on seed nitrogen concentration. We obtained an APHIS permit allowing seed harvest from the first flowering branch. In 2021 at the SoyFACE experimental site, we compared the above-ground biomass, seed weight from the first flowering branch, as well as nitrogen and carbon concentration of seeds from mt2 sprayed twice daily and mt2 without an ethanol treatment. We observed a 35% decrease in chlorophyll concentration at 77 DAP in the ethanol sprayed mutants relative to non-sprayed mutants (Figure 5A). Above-ground biomass and seed weights were not different between the two mutant treatments (Figures 5B and C). However, the nitrogen concentration of the seeds increased in the sprayed low chlorophyll mutants (Figure 5D). We observed a C/N ratio decrease (Figure 5F) and no change in the carbon concentration of the seeds (Figure 5E) between these treatments as well. These results indicate that reducing the chlorophyll concentration of the leaves in the seed filling stages can increase the nitrogen concentration of seeds without a decreasing biomass and seed production under both greenhouse and field conditions when soil N is sufficient.

**Figure 5.**
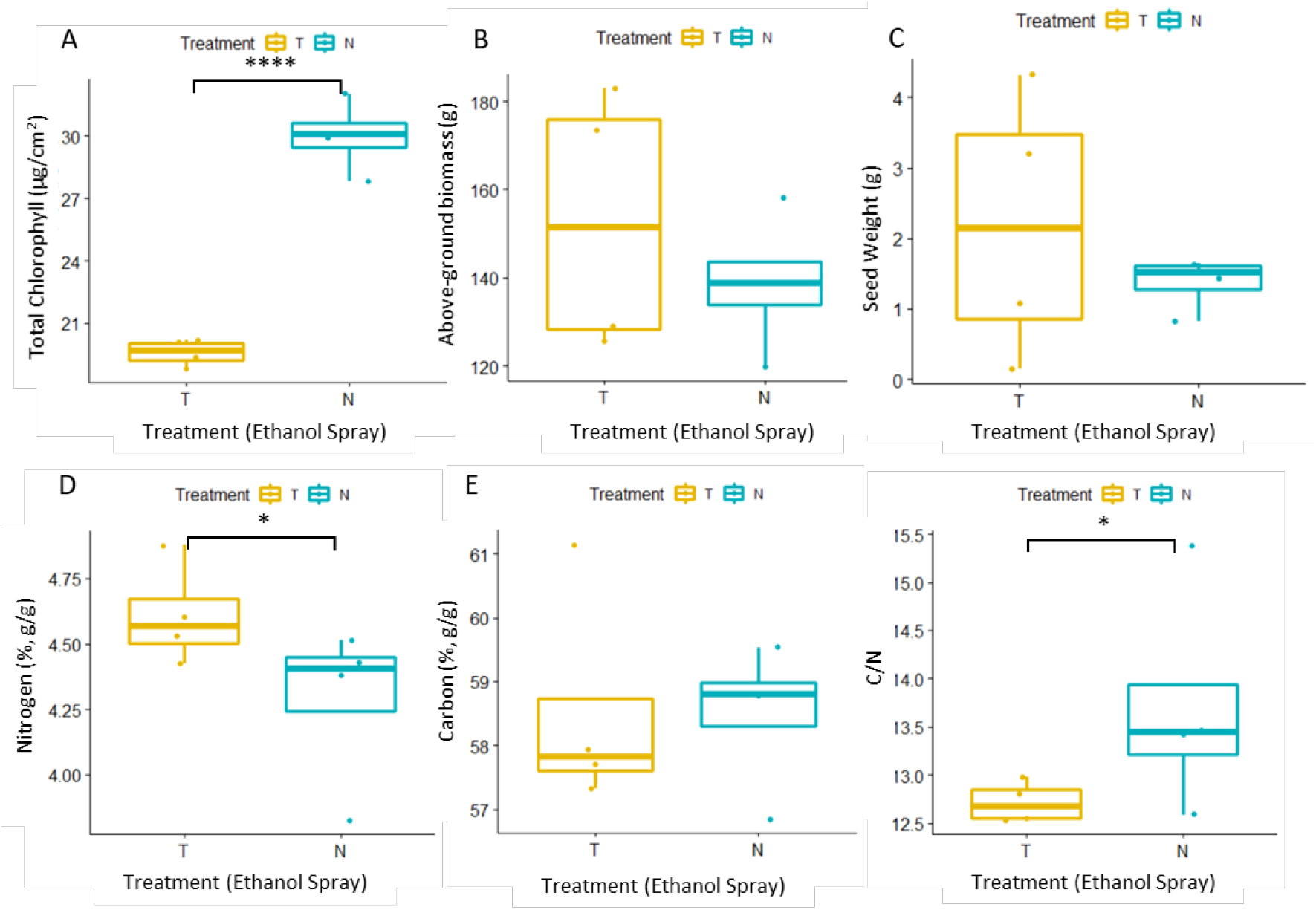
Increased nitrogen concentration and low C/N in seeds of low chlorophyll mutants under field condition. Mutant2 (mt2) tobacco were grown with or without ethanol treatment under field conditions. X-axis shows the ethanol treatment. A. Total chlorophyll level 24 days after treatment (84 DAP,). B. Above ground biomass. C. Seed weight. D. Nitrogen concentration of seed. E. Carbon concentration of seed. F. C/N ratio of seed. The box plots show the median (central line), the lower and upper quartiles (box), and the minimum and maximum values (whiskers). Stars indicate significant difference within treatment when present (*, p<0.1; **, p<0.05; ***, p<0.01,****, p<0.001). The statistical analysis was done using a linear mixed model and type III ANOVA (n= 4 blocks).

## DISCUSSION

### The benefits of inducible gene regulation in specific target periods

In this experiment, we demonstrated the benefits of using an inducible promoter to test our hypothesis that chlorophyll could be significantly reduced without loss of biomass in both greenhouse and field conditions. Although physically inducible promoters, such as heat or light, are perhaps more practical for scalable agriculture than chemically inducible promoters due to environmental issues and operating costs, the ethanol induced Mg chelatase promoter used in this study was more than adequate for proof of concept. In all trials reported in this study regardless of season or location we confirmed reductions of chlorophyll from 50-62% had no effect on biomass relative to neighboring plants with 100% chlorophyll. In addition, we confirmed our prediction in the field and greenhouse that when adequate soil N is available the plant is able to reallocate N not used in chlorophyll production to increase seed protein. We effectively demonstrated that the overproduction of leaf chlorophyll that likely evolved to give these plants a competitive edge is not necessary to maintain expected agricultural yields, and that those resources can effectively be used elsewhere in seed development. The work presented in this study confirms model predictions with empirical evidence and sets the groundwork for numerous future research questions concerning reallocation of chlorophyll N within the plant.

Inducible or time-specific gene regulation may open new opportunities for improving nitrogen-use efficiency in crop plants. Most transgenic plant studies have used constitutive promoters; the 35S promoter has been a favorite choice due to its strong expression often producing a clear phenotype. However, constitutive gene expression of a transgenic trait is not desired in all cases, particularly when targeting essential genes such as Mg-chelatase. In such cases, inducible promoters can provide advantages (Kizis et al., 2001; Selvaraj et al., 2020). For example, an ABA/stress-inducible promoter was found to regulate genes spatially and temporally, improving root architecture (Chen et al., 2015) and drought tolerance (Selvaraj et al., 2020) without yield penalty. Among 41 studies with the goal of improving nitrogen use efficiency (reviewed by Xu et al., 2012), 32 studies used the 35S or ubiquitin promoters to manipulate the expression of genes involved in nitrogen uptake and metabolism. Different timings of fertilizer application have a different effect on yield and seed nitrogen. Yield and seed nitrogen content are highly correlated with nitrogen supply level and availability, particularly at the seed-filling stage (Masclaux-Daubresse and Chardon, 2011). By using the ethanol inducible promoter, we were able to strongly reduce chlorophyll levels once the canopy had closed and plants were producing seeds. Four field trials showed that reducing chlorophyll level did not cause a penalty to above-ground biomass, yield or canopy photosynthesis rate while increasing seed nitrogen concentration (Figures 2-5). This is not possible with a constitutive promoter because reducing Mg-chelatase in early developmental stages leads to staggered growth and sometimes premature death of plants (Supplemental Figure 6), perhaps due to reduced canopy carbon assimilation or pleiotropic effects of down-regulation of *CHLI* (Du et al., 2012; Tomiyama et al., 2014).

Ethanol-inducible promoters have been investigated in greenhouse conditions, but not in field conditions. The expression of the ethanol-inducible promoter is dose dependent for the inducer and sensitive to vapor, which can be used to activate expression (Roslan et al., 2001; Sweetman et al., 2002). The response is very rapid and sustained at a high level; for example, luciferase expression could be detected in the roots of *A. thaliana* seedlings within 1 h of ethanol induction (Roslan et al., 2001). The expression peaks between 3 and 5 days after initial induction (Salter et al., 1998; Garoosi et al., 2005) and maintains a high level via subsequent inductions (Roslan et al., 2001; Schaarschmidt et al., 2004). Similarly, but under field conditions, the current study showed expression of *CHLI* sRNA within 3 hours of ethanol induction, a peak at 3 days, and then reduction of chlorophyll within 5 days (Figures 1E and F). It has been suggested that the ethanol inducible gene expression system is particularly useful for academic research and potentially field use also because ethanol can be broken down by the plant and by microorganisms in the soil (Tomsett et al., 2006). However, the ethanol inducible promoter can be induced unintentionally, because ethanol can be produced in plants under certain conditions. The ethanolic fermentation caused by hypoxia/anoxia converts pyruvate to acetaldehyde, which is later converted to ethanol by ethanol dehydrogenase (Drew, 1997). We observed such an unexpected induction in the Illinois 2019 field experiment, presumably by anoxia (Figure 2C). Between 55-56 DAP, leaves of mt2 in all blocks, including plots without ethanol treatment, turned yellow. After 3 days, chlorophyll production resumed in the untreated mutants. There was 0.3-inch rain when the temperature was 15 °C at the lowest point and 26 °C at the highest point on 52 DAP. It is unclear why only one day of moderate rainfall created an anoxic condition out of all four field trials, but this seems to us the most likely explanation for the induction of the RNAi system in the untreated mutants. Ethanol or acetaldehyde can be produced in some plants naturally, for example in the vascular cambium and xylem sap of trees (MacDonald and Kimmerer, 1991), in large tubers of 16-week-old potato plants (Junker et al., 2004) or in some fruit species (Pesis, 2005). Naturally produced ethanol and acetaldehyde can cause a problem with using an ethanol inducible promoter. We did observe a basal expression of the promoter in mt1 in the greenhouse experiment, but mt2 showed no basal expression in any environment tested (Figures 1 and 2). Tobacco does not naturally produce ethanol and acetaldehyde in normal conditions, but the position of the T-DNA may be at fault in mt1. The location of the T-DNA of mt1 is 1095bp upstream of the transcription start site of the phosphatidylethanolamine binding protein gene, while the T-DNA of mt2 is in the intergenic region (Figure 1C and Supplemental Figure 1). Possibly, the gene cassette for mt1 was expressed by the native promoter of the nearby gene, leading to sRNA expression and reduced chlorophyll in mt1 even without the ethanol signal.

### Lowering leaf chlorophyll content did not negatively impact carbon assimilation or yield, but it increased the seed nitrogen concentration

We evaluated tobacco with reduced chlorophyll over four field seasons and in one greenhouse trial. Once the canopy had closed and seed filling started, we reduced chlorophyll levels by up to 62% without penalty to above-ground biomass or canopy photosynthesis (Figures 2 and 3, Table 1). Overall differences in photosynthesis were not significant despite the reduction of chlorophyll content (Table 1). The minimal changes observed in leaf and canopy photosynthesis are consistent with the lack of reduction in yield for these low chlorophyll plants. These results are in agreement with previous models and experimental studies that predicted or found no differences in yield and photosynthesis between low chlorophyll plants and full green controls (Walker et al., 2018; Song et al., 2017; Gu et al., 2017; Sakowska et al., 2018; Kirst et al., 2017; Slattery et al., 2017). Maintaining yield, biomass, and photosynthesis levels while reducing chlorophyll up to 62% suggests that agricultural plants heavily overinvest in chlorophyll production. As chlorophyll biosynthesis represents a substantial use of nitrogen resources, reducing chlorophyll production could free those nitrogen resources for other uses by the plant that would offer more benefit to agriculture. We showed that reducing chlorophyll in tobacco leaves in later plant developmental stages, after canopy closure and during seed filling, resulted in increased nitrogen concentration in the seeds while maintaining biomass and seed yields (Figures 4 and 5).

Many efforts have been made to maximize nitrogen in the seeds to improve seed quality. The nitrogen content of the seed can be increased when the free amino acid concentrations are increased in the leaf (Caputo et al., 2001) or when the remobilization of leaf nitrogen is accelerated (Taylor et al., 2010; Zhao et al., 2015). Accelerated leaf senescence increases seed protein content (Uauy et al., 2006), while delayed leaf senescence decreases seed protein content (Zhang et al., 2010). Noticeably, delayed leaf senescence increases seed yield due to continuing photosynthetic activity and carbon fixation (Zhang et al., 2010). The timing of leaf senescence controls the mobilization and allocation of carbon and nitrogen resources (Havé et al., 2017). An individual leaf cannot do both carbon assimilation and nitrogen remobilization at the same time, causing the inverse relationship between seed yield and nitrogen content (Havé et al., 2017). Many plants evolved under largely deficient nitrogen conditions compared to modern agriculture and take up nitrogen whenever it is available, to store in Rubisco and other proteins to hoard this normally scarce resource (Denison et al., 2003), even producing more leaves than are required to capture light (Srinivasan et al., 2017). Nitrogen is transferred to seeds only later in development when soil nitrogen is most limiting. Reducing investment in leaves has been proposed to benefit nitrogen use-efficiency under modern agriculture conditions that provide sufficient nitrogen (Denison et al., 2003). Supplying nitrogen during seed filling would be beneficial for crop production (Denison et al., 2003); however, using the nitrogen already stored inside the plants would be more economically advantageous than supplying extra fertilizer during seed filling. Reducing chlorophyll in this study did not initiate senescence, thus leaves were healthy and maintained photosynthetic activity and carbon fixation (Figure 3 and Table 1). The timing of the reallocation of nitrogen resources is critical for the proposed benefit because plants need nitrogen in the leaf for carbon assimilation, so the nitrogen must not be diverted from this purpose too early in development. Lowering the chlorophyll level of the leaf during seed filling is a novel strategy to improve seed quality while maintaining yield.

The effect of reducing chlorophyll on seed nitrogen may vary depending on the availability of nitrogen and plant species. In the greenhouse experiment, we tested five different nitrogen conditions and showed increased nitrogen concentration in the seeds in only two conditions, an adequate nitrogen condition and a slightly lower nitrogen condition (Figure 4). We interpret this observation as the tobacco plants using saved nitrogen in more critical organs such as roots when the nitrogen is very scarce while increasing nitrogen concentration in the seeds only when the demand for nitrogen in critical organs is satisfied. Maintaining root activity during the seed filling stage increased seed nitrogen content (Bogard et al., 2010). Further study is required to investigate the root system as the root is both a sink and source tissue of nitrogen. Controlling nitrogen remobilization has a different effect depending on the species. Each species has a different portion of seed nitrogen that comes from the remobilization of nitrogen stored in roots and shoots before flowering: wheat (Triticum aestivum L.) is 60–95% (Palta and Fillery, 1995); maize (Zea mays L.) is 45–65% (Rajcan and Tollenaar, 1999); rice (Oryza sativa L.) is 70–90%. Tobacco (cv. Petite Havana) like other plants also remobilizes nitrogen from leaves to sink reproductive tissues when flowering. In this study, the chlorophyll levels of wild-type and mutants kept increasing right before flowering, then decreased during seed filling and bounced back within 5-6 days (Figures 2C and E). Demands for nitrogen can stimulate organic nitrogen remobilization that is mediated by autophagy and vacuolar proteases (Tegeder et al., 2014). Nitrogen-related macromolecules such as Rubisco and the chlorophyll embedded light-harvesting complex are broken down into amino acids, peptides and ureides, then transferred to nitrogen-demanding organs by various transporters (Tegeder et al., 2014).

It is not elucidated yet where low-chlorophyll tobacco plants store saved nitrogen from reducing chlorophyll and how the targeting and regulation of the nitrogen reallocation is managed. Perhaps, reducing chlorophyll via down-regulation of Mg-chelatase leads to a breakdown of excess light-harvesting complexes, then the organic nitrogen is remobilized to sink tissues such as seeds. Analysis of the form of nitrogen in harvested leaves from one field season, IL2019, showed both organic and inorganic nitrogen concentrations were higher in reduced chlorophyll leaves subjected to the once-a-day ethanol treatment (Supplemental Figure 5). Saved nitrogen can be remobilized in both organic and inorganic forms in plants. Inorganic nitrogen can be remobilized during leaf senescence depending on the size of the inorganic pool stored in leaves before senescence, although remobilized organic nitrogen has a larger effect on seed nitrogen content (Havé et al., 2017). We speculate that reducing chlorophyll increases the pool size of the inorganic nitrogen in leaves as nitrate content in low-chlorophyll leaves is significantly higher (Supplemental Figure 5). Another possible explanation is that lowering chlorophyll in leaves reduces nitrogen demands there, leading to seeds being the sole sink tissue thus, nitrogen from the soil and root is transferred only to seeds. Further study will investigate how lowering chlorophyll distributes nitrogen by tracking nitrogen isotopes given in different treatments and different developmental stages.

It is tempting to propose that a reduction in chlorophyll production can maintain the quality of seeds in future climate environments where the elevated CO_2_ concentration accelerates carbon assimilation and increases seed carbon to nitrogen ratios in non-leguminous species (Myers et al., 2014). Increased carbon assimilation changes the stoichiometric balance of nutrients in plants by increasing carboxylation, but not the absorption of minerals (Loladze, 2002). A meta-analysis of free-air carbon dioxide enrichment (FACE) experiments showed that wheat seed protein was reduced by 6% and rice seed protein by 8% under the elevated CO_2_ conditions that also led to increased photosynthesis (Myers et al., 2014), which would disturb the carbon to nitrogen (C/N) balance by producing more carbon (McGrath and Lobell, 2013). Our work here demonstrates that reducing chlorophyll by about 60% in the leaves increases the nitrogen concentration of the seeds by 4-17% depending upon the amount of nitrogen fertilization (Figures 4 and 5). Therefore, reducing chlorophyll could maintain the C/N ratio by increasing nitrogen to match the increased carbon from improved photosynthesis. The balance of these two nutrients, markers of seed quality, could be maintained with no penalty on biomass or seed production using this method. A further FACE experiment will illuminate the benefit of reducing chlorophyll in climate change.

In summary, to understand the benefits of reducing chlorophyll in later developmental stages, we created an inducible RNAi tobacco mutant that expresses Mg-chelatase subunit I (*CHLI*) sRNA within 3 hours of induction and reduces chlorophyll within 5 days in a field condition. Once the plants had grown enough and the canopy closed, we reduced chlorophyll levels by up to 62% without penalty to above-ground biomass or canopy photosynthesis. Chlorophyll reduction of the leaf in the seed filling stages increased nitrogen concentration in the seeds, while biomass and seed yields were maintained. On top of the breeding efforts to maximize yield and seed quality, we suggest the time-specific reduction of chlorophyll as a niche strategy to decouple the inverse relationship between yield and seed nitrogen.

## METHODS

### Cloning and transformation

The constructs were generated using Golden Gate cloning (Engler et al., 2008; Engler et al., 2009). Level 0 constructs including *alcR* regulon (EC27885), *alcA* promoter (EC27886), *GA20* intron (EC27888), and *CHLI* RNAi (EC27891) were newly synthesized based on the sequence used in Chen et al. (2003). The assembled construct is described in the Results section (Figure 1B). *N. tabacum* cv. Petit Havana was genetically transformed using *Agrobacterium tumefaciens* mediated transformation (Gallois and Marinho, 1995). Twenty-two independent T0 transformations were generated to produce T1 progeny. T-DNA copy number was determined on T1 plants through copy number quantitative polymerase chain reaction (qRT-PCR) analysis (iDNA Genetics, Norwich, UK) (Supplemental Table 3). From these results, two events were selected based on the leaf color after ethanol treatment in the greenhouse. Then, the two selected lines were selfed and produced T2 progeny with copy numbers confirmed for the homozygous single insertion (Supplemental Table 3). T3 progeny carrying a homozygous single insertion was used in the experiments.

### T-DNA location

Genomic DNA was extracted from young leaves as described in Cho et al., 2019. The locations of T-DNA in two mutants were revealed by targeted locus amplification performed by Cergentis B.V. (Utrecht, Netherlands) (De Vree et al., 2014). The location of T-DNA in mt1 was confirmed by PCR using a primer set; the forward primer resides on the native tobacco genome near the breakpoint and the reverse primer resides on the T-DNA. The location of T-DNA in mt2 was confirmed by using the GenomeWalker kit (Clontech Laboratories, Mountain View, CA) following the manufacturer’s recommended protocols. Both locations were confirmed by PCR using unique primer sets. All primer sequences are described in Supplemental Table 4.

### Greenhouse and field conditions

In all greenhouse trials despite the year and location, plants were grown as described in South et al. (2019). T3 seeds were germinated using Berger 6 mix (Berger, Saint-Modeste, QC, Canada). Thirteen days after germination, seedlings were transferred to 6L pots (600C, Hummert International, Earth City, MO, USA) with Berger 6 mix supplemented with slow-release fertilizer (Osmocote Plus 15/9/12, The Scotts Company LLC, Marysville, OH, USA). We added 0, 0.2, 0.4, and 0.8 grams of urea to Sandy Loam Mix (sand:soil, 2:1) instead of the Osmocote for the nitrogen experiments. Plants were cultivated in greenhouse conditions; daytime 26 °C– 28 °C, nighttime 23 °C– 25 °C, 16 h photoperiod with natural light supplemented under low light induced by cloud cover with high-pressure sodium light bulbs, giving 300–800 μmol m^−2^ s^−1^ from the pot level to the top of the plant.

In all field trials despite the year and location, each field was prepared in a similar fashion each year, as described by Kromdijk et al. (2016). Plants were grown using a method broadly similar to commercially grown tobacco. Flowers were removed, collected, and dried to add into the above-ground biomass. Above-ground biomass was measured after drying in an oven (58-60 °C) for at least 2 weeks to constant weight before biomass measurements. All trials had randomized complete block designs with different genotypes and treatments as described in the Results section and Supplemental Figure 7.

### Ethanol treatment

For the root drenching, 100 ml of 1% (v/v) ethanol was applied to the roots (Figure 1D). For all leaf spraying, fresh 2% (v/v) ethanol was prepared every morning in both greenhouse and field experiments. A hand-held sprayer (Chapin 26021XP 2-Gallon ProSeries Poly Sprayer, Batavia, NY) was used until sun leaves (4-5th leaf from the top, exposed to full sunlight) were fully soaked with the solution. Spray time was between 7-9 am, or after 5 pm to avoid full sunlight.

### Field experiments for above-ground biomass and canopy photosynthesis

#### Illinois 2019

To evaluate the impact of the (*CHLI)* mutation induced by the ethanol spray on chlorophyll level and above-ground biomass, we used three genotypes (wild-type, mutant1, and mutant2) each with three spray treatments; spray once a day (-1), spray twice a day (-2), or no spray (−0). Plants were planted in pots containing soil mix on 31 May 2019 and grown for 12 days before transfer to floating trays in the greenhouse. Plants were transplanted at the University of Illinois Energy Farm field station (40.06° N, 88.21° W, Urbana, IL, USA) on 27 June 2019 (27 DAP). Each plot consisted of 6 × 6 plants spaced 30 cm apart. The ethanol treatment began on 18 July 2019 (48 DAP). The whole canopy was maintained until the harvest. Seven internal plants per plot were harvested and dried on 20 Aug 2019 (81 DAP) to measure aboveground biomass. All border and remaining internal plants were not used.

#### Puerto Rico 2019

To evaluate the impact of reduced chlorophyll levels on the above-ground biomass, in this experiment we used three genotypes (wild-type, mutant1, and mutant2), each with one spray treatment, twice daily. Plants were germinated in pots containing Berger 6 soil mix on 1 October 2019 and grown for 17 days before transfer to floating trays. Plants were transplanted at the IL Crop Field Station (18.00° N, 66.51° W, Ponce, Puerto Rico) on 11 November 2019 (41 DAP). Each plot consisted of 5 × 5 plants spaced 22 cm apart. The ethanol treatment began on 25 November 2019 (55 DAP). The whole canopy was maintained until the harvest. Above-ground biomass of all nine internal plants per plot was harvested and dried on 14 December 2019 (74 DAP). Border plant biomass was not measured.

#### Illinois 2020

Since trials in 2019 demonstrated that the genotype mutant1 did not respond to sprayed ethanol, only two genotypes (wild-type and mutant2) were used in 2020 to evaluate the impact of reduced chlorophyll on aboveground biomass. Two genotypes within one treatment (WT-2 and mt2-2) and two treatments within one genotype (mt2-2 and mt2-0) were compared. All were planted in pots containing soil mix on 16 May 2020 and grown for 16 days before transfer to floating trays. Plants were transplanted at the University of Illinois Energy Farm field station (40.11°N, 88.21°W, Urbana, IL, USA) on 11 June 2020 (26 DAP). Each plot consisted of 5 × 5 plants spaced 22 cm apart. Due to a hail storm on 11 July 2020 (56 DAP) that created variable canopy damage, we coppiced all plants 12 cm above the ground on 14 July 2020 (59 DAP) and let the plants regrow (Supplemental Figure 8). The ethanol treatment began on 27 July 2020 (72 DAP). The whole canopy was maintained until the harvest. The nine internal plants per plot were harvested and dried on 2 September 2020 (109 DAP). Border plant biomass was not measured.

### Greenhouse and field experiments for above-ground biomass and seed analysis

#### Greenhouse 2020

To test the theory that freed N from reduced chlorophyll could be used for seed protein we conducted a greenhouse experiment with wildtype and mutant2 plants with four N treatments. Above-ground biomass, seed yield, and seed composition were measured on plants with one ethanol spray treatment, twice daily. The treatment began at 60 DAP, when flowering began (followed shortly by seed filling). Above-ground plant material was collected at 101 DAP to measure dry biomass, seed yield, and seed tissue composition.

#### SoyFACE 2021

To further elucidate the relationship between reduced chlorophyll and seed composition we conducted a field experiment comparing above-ground biomass, seed yield, and seed composition in mutant2 plants with and without a sprayed ethanol treatment (twice a day). Plants were planted in pots containing soil mix on 2 July 2021 and grown for 16 days before transferring to floating trays. Plants were transplanted at the SoyFACE field station (40.04°N, 88.23°W, Urbana, IL, USA) on 5 August 2021 (34 DAP). Each plot consisted of 4 plants spaced 22 cm apart. The first branch of the flower was covered with a silk cloth to collect seeds. The ethanol treatment began on 31 August 2021 (60 DAP). All above-ground plant material were harvested and dried on 12 October 2021 (102 DAP) to measure dry biomass, seed yield, and seed tissue composition.

### Chlorophyll measurement

SPAD measurements for chlorophyll content were taken at the same spot on the leaf as the gas exchange measurements for the photosynthetic response curves. To convert SPAD readings to chlorophyll content using a linear function (Supplemental Figure 3), we collected leaf disks 1 cm in diameter (the same area measured by SPAD) in one field (IL2019) and greenhouse experiment, which were frozen in liquid nitrogen for at least 10 minutes and stored in the freezer (−80 °C) until they were lyophilized (Benchtop Freeze Dryers, Labconco Co. Kansas City, MO). Chlorophyll content was then determined using 100% ethanol extraction (Ritchie, 2006) and analyzed with spectrophotometer (BioTek PowerWave Microplate Reader, Agilent, Santa Clara, CA). For other experiments, chlorophyll content was measured by SPAD (Soil Plant Analysis Development; Chlorophyll Meter SPAD-502 Plus; Konica Minolta, Japan) on a sun leaf. Three (before ethanol treatments) to nine (after ethanol treatments) plants per plot were measured in all field experiments, and every plant was measured in the greenhouse.

### sRNA extraction and next generation sequencing analysis

Leaf disks 1 cm in diameter were collected from the same leaf at multiple time points: 0, 1, 3, 6, 24, 48, 72, 96, and 336 hours after the first ethanol spray. Leaf discs were immediately frozen in liquid nitrogen for at least 10 minutes and stored in the freezer (−80 °C) until they were lyophilized. Total RNA was isolated from freeze-dried leaf disks using phenol and chloroform extractions (Wang and Vodkin, 1994) and precipitated with ethanol but without lithium chloride to preserve sRNAs (Cho et al., 2013; Cho et al., 2017). The small RNA libraries were prepared using the NEBNext Small RNA Sample Prep kit (New England Biolabs, Ipswich, MA). High throughput sequencing was performed with NovaSeq-6000 (Illumina, San Diego, CA) by the Keck Center (University of Illinois, Urbana, IL) using Illumina protocols. Generally, a total of 8 to 10 million reads were obtained from these deep-sequencing libraries. Adapter-trimmed sequences were aligned to the tobacco genome (Edwards et al., 2017) and quantified by using the program Salmon (Patro et al., 2017).

### *A/C_i_* response curves

The response of *A* to *C_i_* in randomly chosen sun and shade leaves was measured at a saturating light intensity of 2000 μmol quanta mol^−2^ s^−1^ by using a portable infrared gas analyzer (LI-COR 6800; LI-COR, Lincoln, NE). Illumination was provided by a red–blue fluorometer light source attached to the leaf cuvette. Measurements of *A* were started at the ambient CO_2_ concentration (*C_a_*) of 400 μmol mol^−1^, before *C_a_* was decreased stepwise to a lowest concentration of 0 μmol mol^−1^ and then increased stepwise to an upper concentration of 1,300 μmol mol^−1^ (400, 300, 200, 100, 50, 0, 400, 400, 600, 800, 1000, and 1300 μmol mol^−1^). All parameters were calculated from *A* versus *C_i_* by fitting the data to a non-rectangular hyperbola. The maximum carboxylation efficiency (*CE*), the CO_2_ compensation point (*Γ*), the CO_2_ saturated rate of *A_ne_*_t_ (*A_sat_*), and the empirical curvature factor for the *A/C_i_* curves (*ω*) were calculated from *A/C_i_* measurements.

### *A/Q_abs_* response curves

Photosynthesis as a function of light (*A/Q_abs_* response curves) in randomly chosen sun and shade leaves was measured under the same cuvette conditions as the *A/C_i_* curves mentioned above at an ambient CO_2_ concentration (*C_a_*) of 400 μmol mol^−1^ by using a portable infrared gas analyzer (LI-COR 6800; LI-COR, Lincoln, NE). The leaves were initially stabilized at a saturating irradiance of 2000 μmol quanta m^−2^ s^−1^, after which *A* and *g_s_* were measured at the following light levels: 2000, 1500, 1000, 750, 500, 200, 180, 160, 140, 120, 100, 80, 60, 40, 20, and 0 μmol m^−2^ s^−1^. The measurements were recorded after *A* reached a new steady state (1–2 min) and before *g_s_* stabilized to the new light levels. All parameters were calculated from *A* versus absorbed PPFD by fitting the data to a non-rectangular hyperbola. A quadratic relationship between leaf light absorptance and chlorophyll level (SPAD) was determined using an integrating sphere (Jaz Spectroclip, Ocean Optics, Duiven, Netherlands) as described in Walker et al. (2018) (Supplmental Figure 4D). The maximum quantum yield of CO_2_ assimilation (*ΦCO_2_*), respiration rate (*R*), the light saturated rate of net CO_2_ assimilation (*A_sa_*_t_), the empirical curvature factor (*θ*), and the maximal photochemical yield of photosystem II obtained at low light (*ΦPSII(LL)*) were calculated from *A/Q_abs_* curves.

### *A’* calculations

A simple three leaf-layer concept was used to model canopy photosynthetic rates (Supplemental Figure 4A) simulating an LAI of 3 similar to measurements from two field trials (Supplemental Figure 5). Modeled parameters included diurnal photosynthetic photon flux density (PPFD) incident on the uppermost leaf layer (Supplemental Figure 4B), linear relationships between transmittance and reflectance (Supplemental Figure 4C), a quadratic relationship between leaf light absorptance, chlorophyll level (SPAD) (Supplemental Figure 4D) and the photosynthesis parameters from the light response curves (*A/Q_abs_* curves; Table 1). These parameters were used to calculate light absorbed and carbon assimilated throughout a 24-hour period; the carbon assimilated in each leaf layer was summed to calculate the total net carbon assimilated by the canopy (*A’*).

### Seed composition analysis

Dried seeds were homogenized and ground into a powder with a ball mill (Geno Grinder 2010; BT&C Lebanon New Jersey, USA). Ground material was weighed into tin capsules for C and N analysis, and combusted with an elemental analyzer (Costech 4010CHNS Analyzer, Costech Analytical Technologies Inc. Valencia, California, USA). Acetanalide and Apple Leaves (National Institute of Science and Technology, Gaithersburg Maryland, USA) were used as standards as in Masters et al. (2016).

### Leaf composition analysis and LAI measurement

Dried leaves from the harvested plants in the IL2019 field trial were collected and analyzed to investigate the form of nitrogen. Tissue samples were sent to a commercial lab (Midwest Laboratories, Omaha, NE) where the samples were ground, homogenized, and analyzed for total N (Dumas method with a KECI FP428 nitrogen analyzer, AOAC method 968.06), nitrate N (cadmium reduction automated FIA determination, AOAC 968.07), ammonia N (distillation method for the determination of ammonia nitrogen, AOAC 920.03) and organic N (Kjeldahl method, AOAC 2001.11). The same ground samples were analyzed by elemental analyzer (see above for method) to calculate C/N ratio. LAI was determined for three (IL2020) to four (IL2019) plants in each plot using a leaf area meter (LI-3100, LI-COR, Lincoln, NE, USA).

### Statistical analysis

The statistical analyses were done by using SAS (SAS Institute, Cary, NC) and R (version 4.0.3). The biomass, photosynthesis, and chlorophyll levels from field trials were analyzed in a mixed model analysis of variance (ANOVA) and posthoc Tukey test (α=0.05). Block was considered as the random effect for all analyses, while genotypes and spray frequency (IL2019), genotype (PR2019), and genotype-ethanol spray (IL2020) were considered as the fixed effects. For the greenhouse experiment, ANOVA was used. If data were not normally distributed, non-parametric analysis (Wilcoxon rank test) was conducted using R.

## DATA AVAILABILITY

All study data are included in the article and/or SI Appendix.

## ACKNOWLEDGMENTS

We thank D. Drag, B. Harbaugh, B. Thompson, R. Edquilang and R. Metallo for plant care and management in the greenhouse and field studies; and M. Kim, D. Chernyakhovskiy, D. Nguyen for assistance during laboratory and field work. We thank M. Balasubmaranian for tobacco transformation. We thank S. Burgess for critical review of the manuscript. Funding: This work is supported by the research project Realizing Increased Photosynthetic Efficiency (RIPE) that is funded by the Bill & Melinda Gates Foundation, Foundation for Food and Agriculture Research, and U.K. Foreign, Commonwealth & Development Office under grant no. OPP1172157. This work is licensed under a Creative Commons Attribution 4.0 International (CC BY 4.0) license, which permits unrestricted use, distribution, and reproduction in any medium, provided the original work is properly cited. To view a copy of this license, visit https://creativecommons.org/licenses/by/4.0/. This license does not apply to figures/photos/artwork or other content included in the article that is credited to a third party; obtain authorization from the rights holder before using such material.

---

**Supplemental Figure 1.**
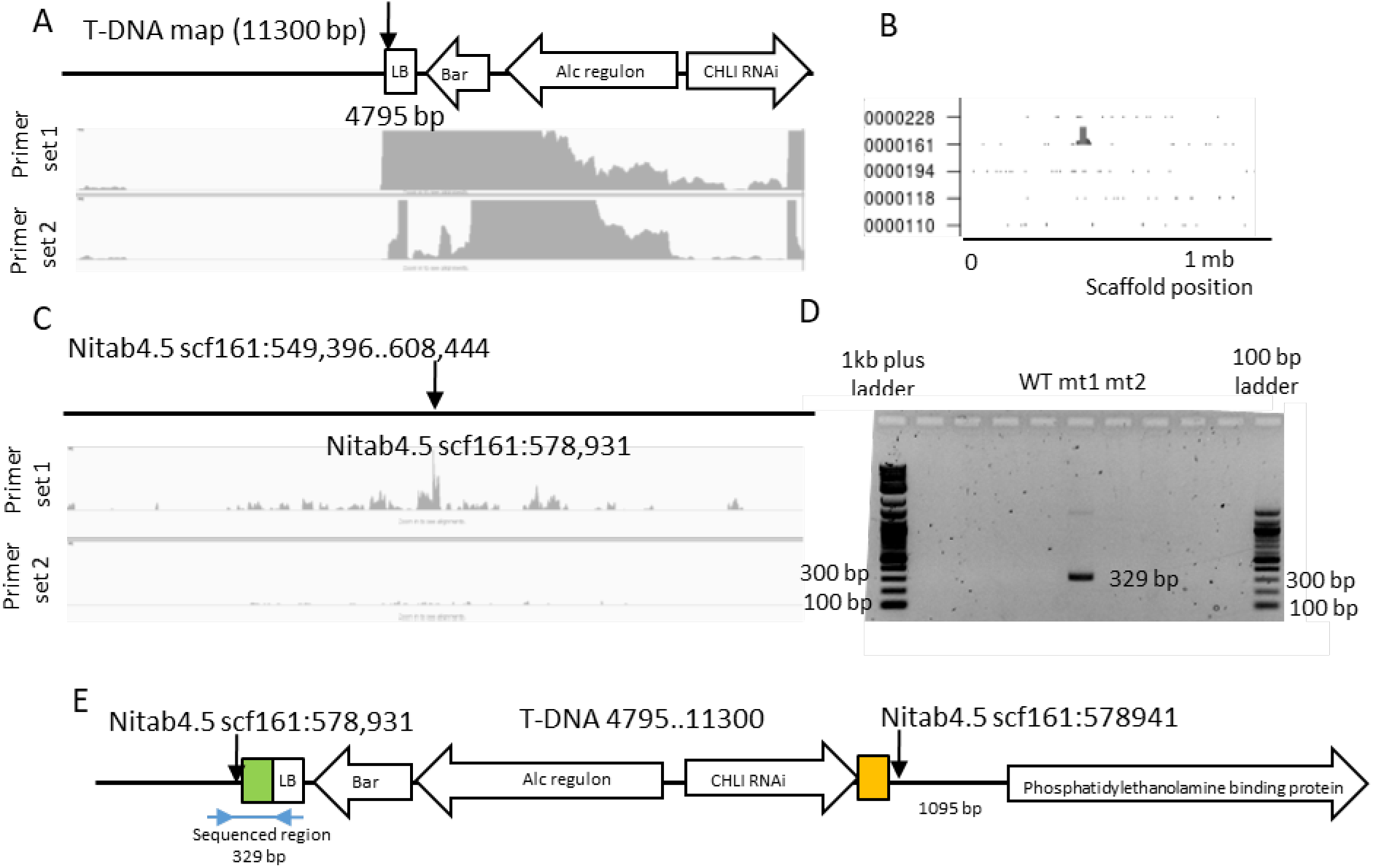
T-DNA location of mt1. A. Transgene sequencing coverage: NGS sequencing coverage across the TG with primer set 1 (top) and primer set 2 (bottom). Black arrows indicate primer location. Y-axis is limited to 100x. B. Whole genome coverage plot: TLA (Targeted locus amplification) sequence coverage across the top 50 scaffolds of the tobacco genome using primer set 1. The scaffolds are indicated on the y-axis, the scaffold position on the x-axis. No genome coverage was observed with primer set 2 (data not shown). Scaffold 161 shows the coverage of TLA sequence. C. TLA sequence coverage across the TG integration locus; tobacco scaffold 161:549,396-608,444. Sequence coverage (in grey) generated with primer set 1 (top) and primer set 2 (bottom). The black arrow indicates the breakpoint sequences (scf161:578931) identified with primer set 1. Y-axis is limited to 50x. D. Amplicon for the chimeric region. The forward primer resides on the native tobacco genome near the breakpoint and the reverse primer resides on the T-DNA. Amplicon is found only in mt1, confirms the location of T-DNA in mt1. E. Schematic figure of the chimeric tobacco genome sequence. Tobacco genome (scf161:549,396..555,535) linked to left border (4795 bp) sequence of T-DNA. Green (61 bp) and orange boxes represent unknown sequence between native tobacco genome and T-DNA.

**Supplemental Figure 2.**
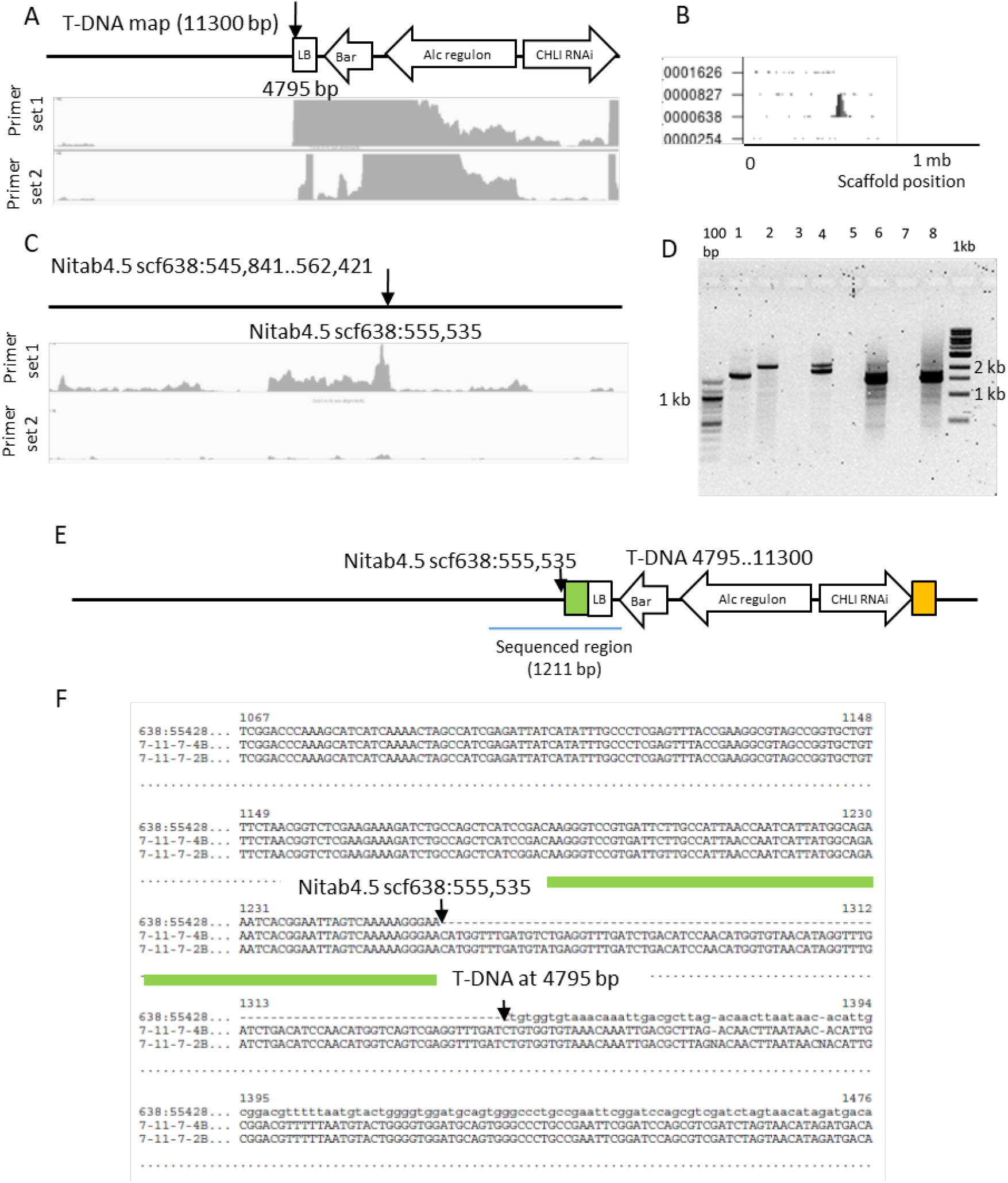
T-DNA location of mt2. A. Transgene sequencing coverage: next generation sequencing coverage across the transgene with primer set 1 and 2. Y-axis is limited to 100x. Black arrow indicates the break point at the left border of T-DNA construct (4795 bp). B. Whole genome coverage plot: TLA (Targeted locus amplification) sequence coverage across the top 50 scaffolds of the tobacco genome using primer set 1. The scaffolds are indicated on the y-axis, the scaffold position on the x-axis. No genome coverage was observed with primer set 2 (data not shown). Scaffold 638 shows the coverage of TLA sequence. C. TLA sequence coverage across the transgene integration locus; tobacco scaffold 638:545,841-562,421. Sequence coverage (in grey) generated with primer set 1. The black arrow indicates the breakpoint sequences (scf638:555,535) identified with primer set 1. Y-axis is limited to 50x. D-E. The sequence of the chimeric region of T-DNA integration site confirmed by using Genome Walker kit (Takara). D. Gel lane 1: digested by DraI, 2: EcoRV, 3: PvuII, 4: StuI, 5 and 7: negative controls. 6 and 8: positive controls. E. Schematic figure of the chimeric tobacco genome sequence. Tobacco genome scf638:554,535 – 555,535 linked to left border sequence of T-DNA. Green (90 bp) and orange boxes represent unknown sequence between native tobacco genome and T-DNA. F. The sequences of the amplicons (7-11-7-4B and 7-11-7-2B) are aligned to the chimeric genome sequence (tobacco genome scf638:554,535 – 555,535 linked to left border sequence of T-DNA).

**Supplemental Figure 3.**
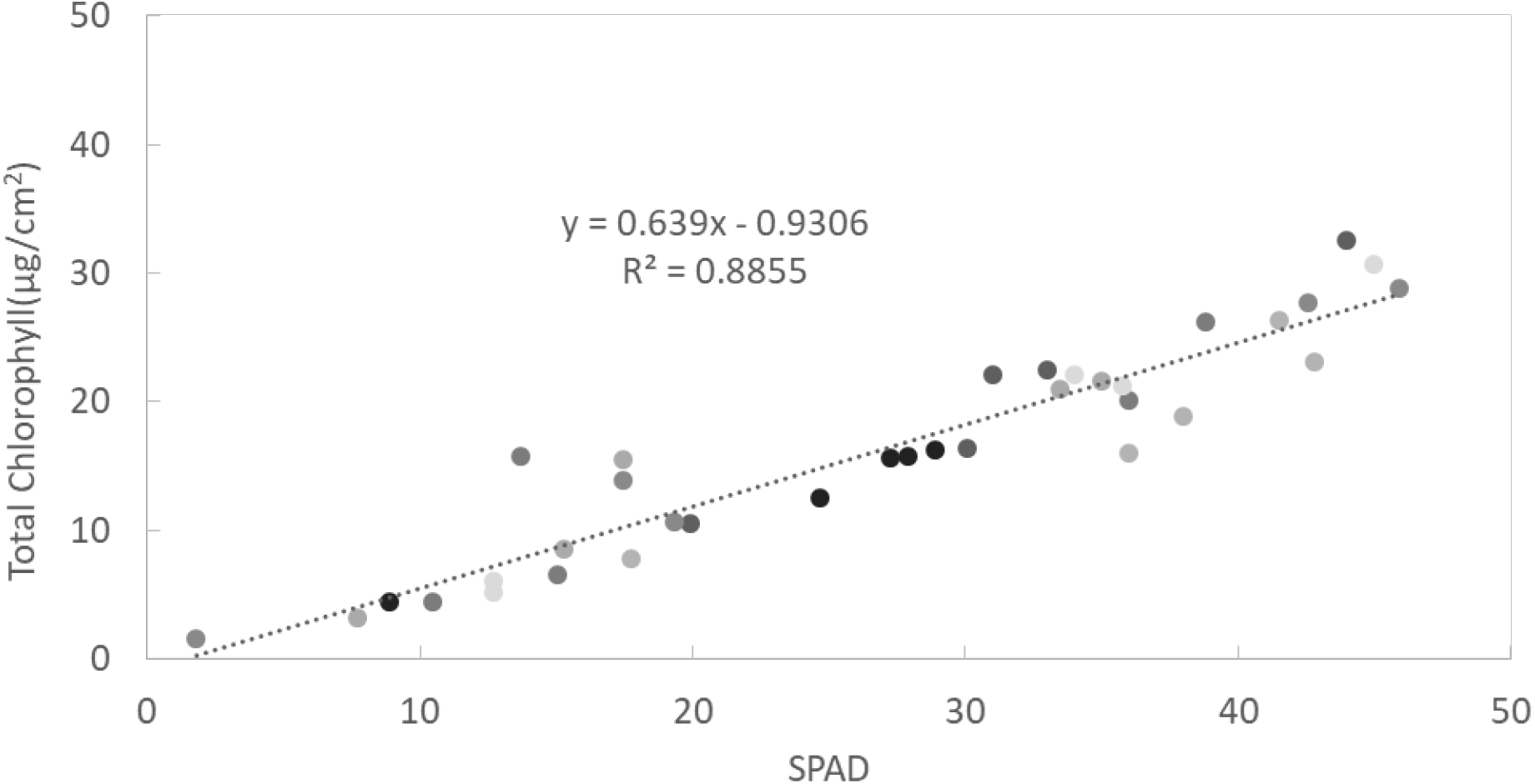
Relationship between SPAD and total chlorophyll content in tobacco. Leaf disks 1 cm in diameter (the same area measured by SPAD) were collected in the field and greenhouse, then frozen in liquid nitrogen for at least 10 minutes and stored in the freezer (−80°C) until they were lyophilized. Chlorophyll content was determined using ethanol extraction. The relationship used to convert SPAD readings to chlorophyll amount was calculated using a linear function.

**Supplemental Figure 4.**
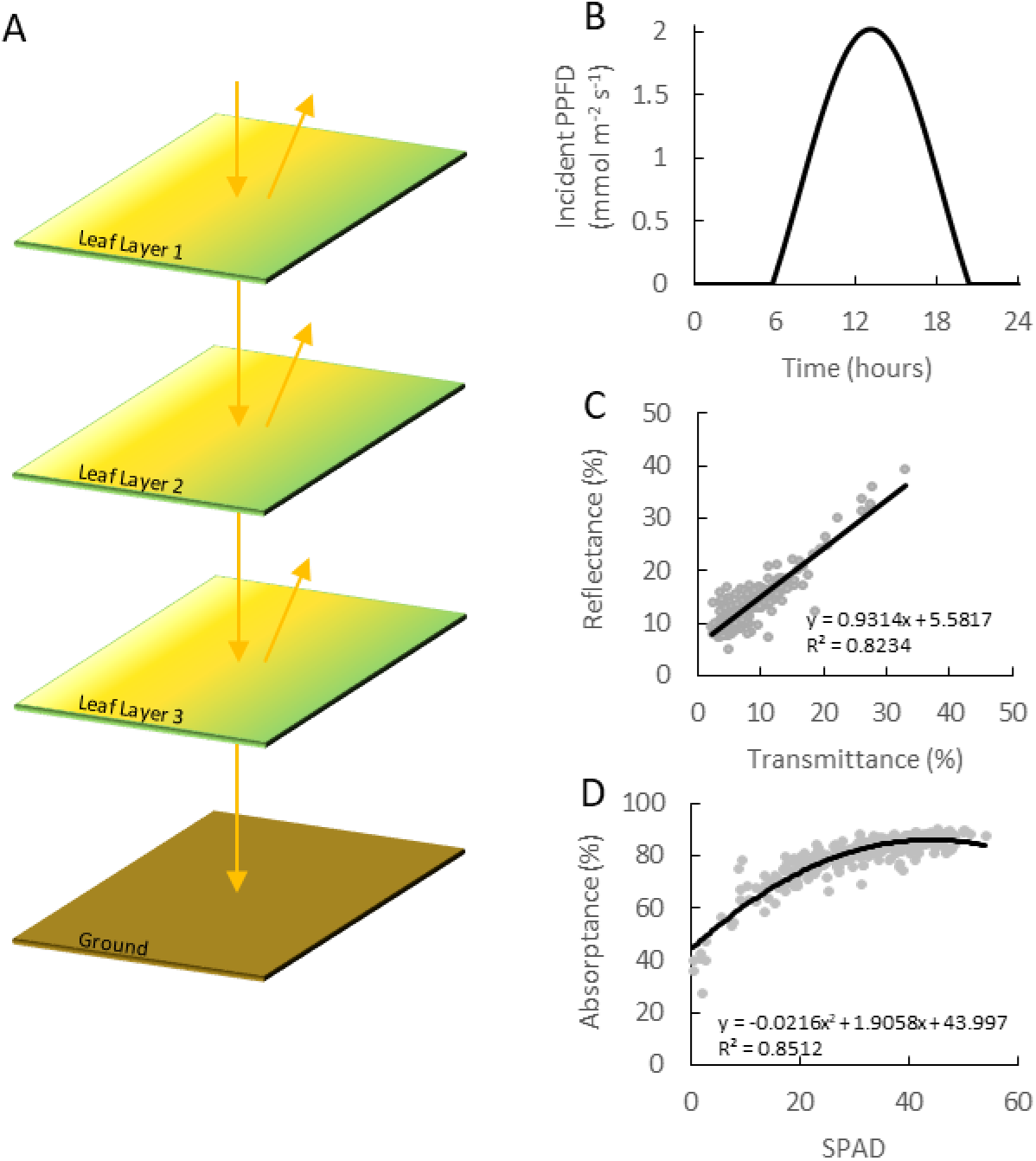
Total net carbon assimilation of the canopy (A’) is calculated by using an artificial canopy model. A. Three-leaf concept used to model canopy photosynthetic rates. B. Diurnal photosynthetic photon flux density (PPFD) incident on the uppermost leaf layer (Leaf Layer 1). C. Linear model fits of reflectance to transmittance. D. Empirical relationship between leaf light absorptance and SPAD value.

**Supplemental Figure 5.**
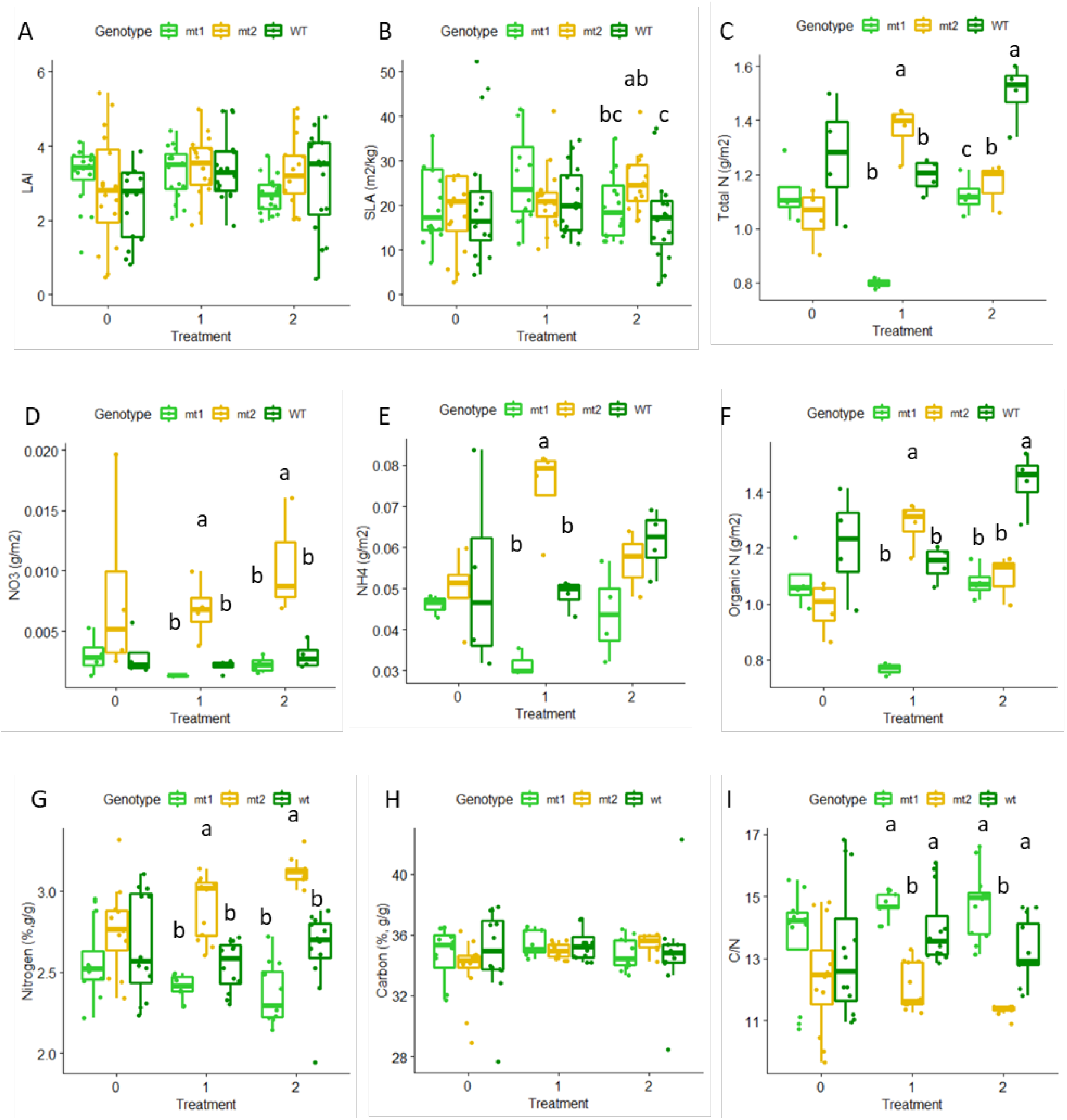
Leaf composition analysis IL 2019 field experiments. A. LAI B. SLA C. Total nitrogen content (area-based). D. Nitrate content (area-based). E. Ammonia content (area-based). F. Organic nitrogen content (area-based). G. Nitrogen content (weight-based). H. Carbon content (weight-based). I. Carbon to nitrogen ratio. A and B. Four plants collected from each plot to measure leaf area and weight. C-H. Dried leaves from each plot collected and ground to homogenize. The box plots show the median (central line), the lower and upper quartiles (box) and the minimum and maximum values (whiskers). The statistical analysis was done using ANOVA and post-hoc Tukey test. Letters indicate significant differences within treatment when present.

**Supplemental Figure 6.**
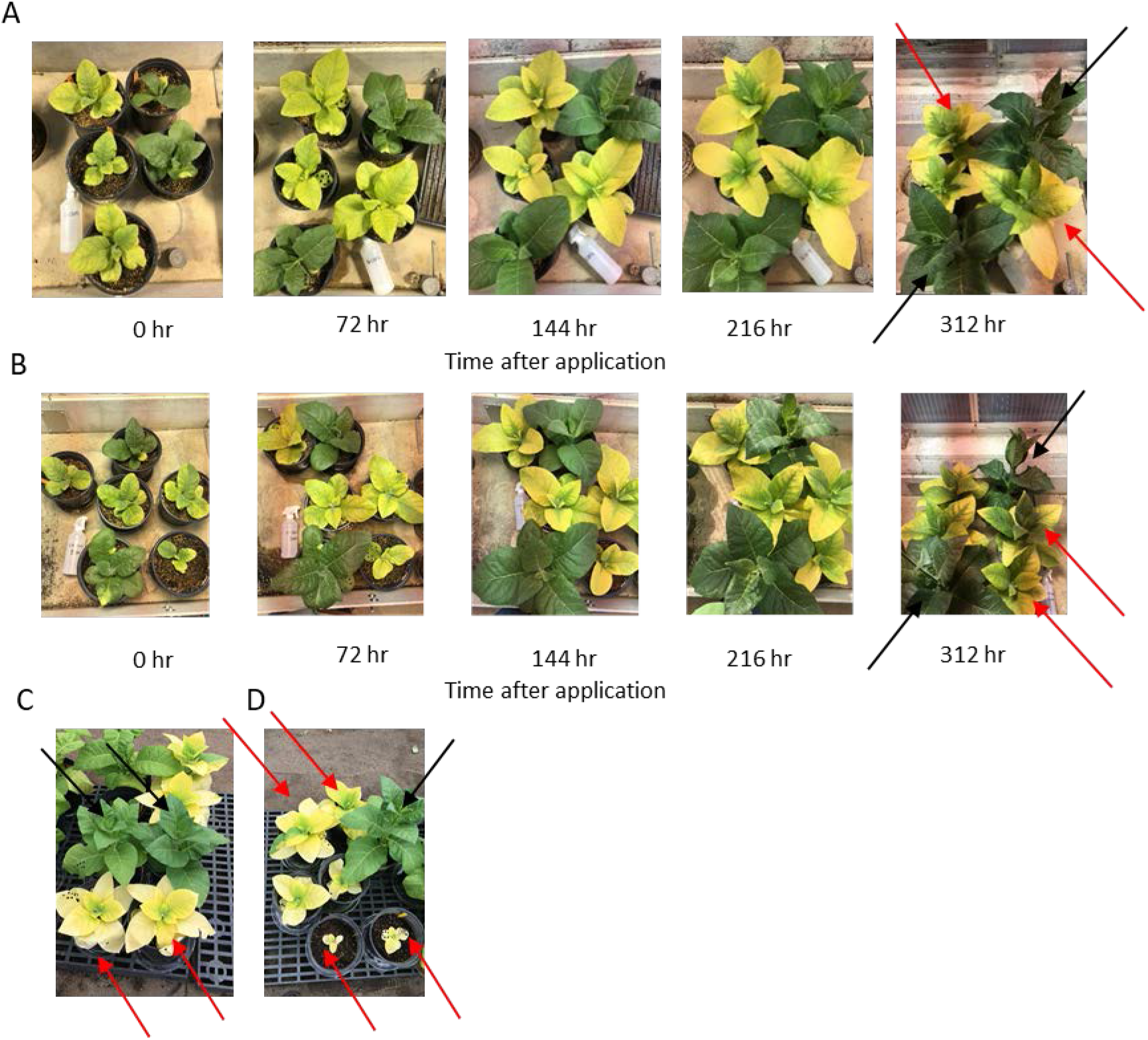
Early induction of chlorophyll down-regulation inhibits the growth of mutants in the greenhouse. A. Change of leaf color after 1% ethanol spray in tobacco. Red arrows indicate affected mutants showing retardation of growth compared to the dark green azygous (black arrow). B. Change of leaf color after 0.5% ethanol spray. C-D. Earlier induction (beginning with germination) causes significant growth retardation and premature death of tobacco (Samsun).

**Supplemental Figure 7.**
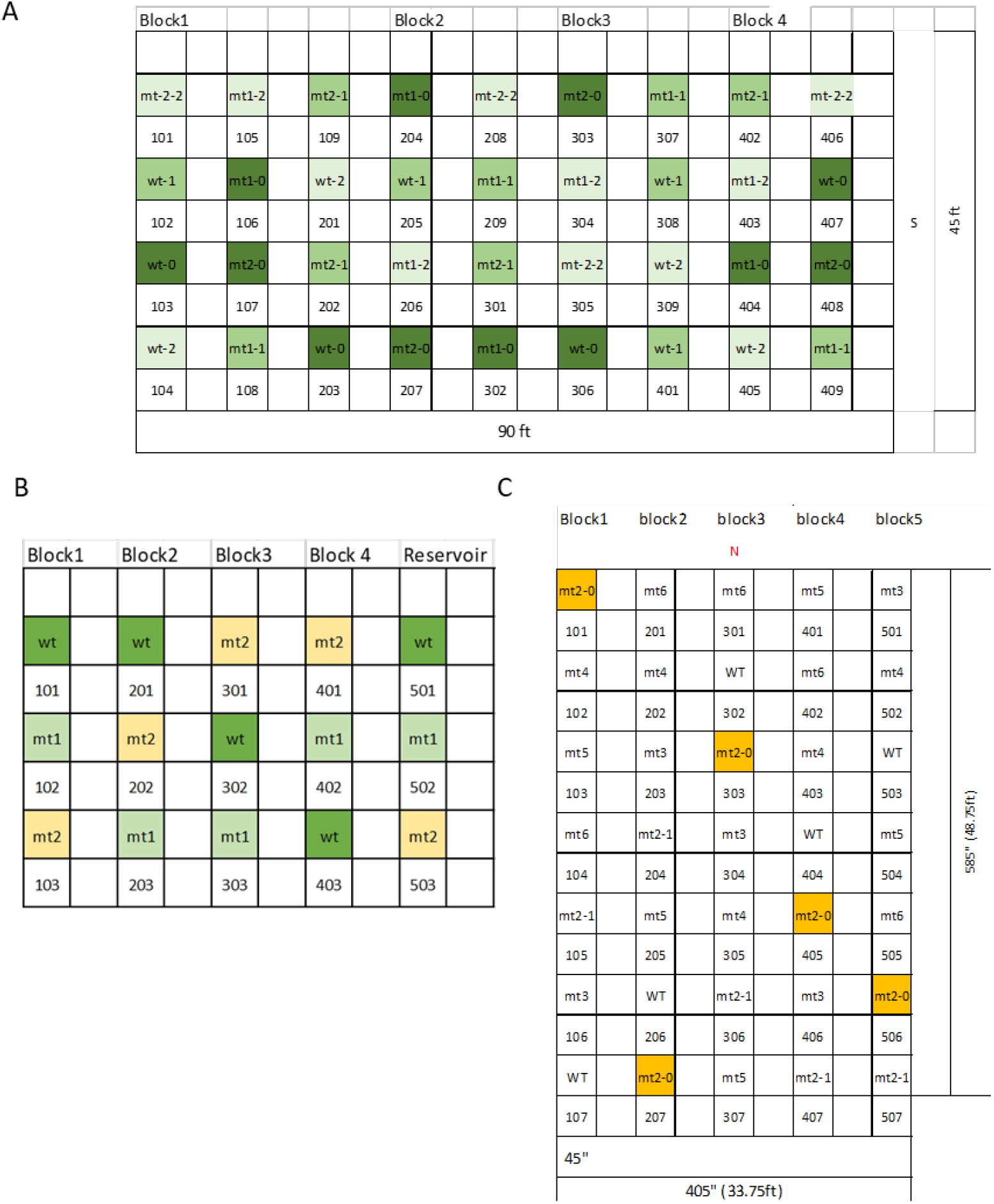
Schematic representation of field trials. A. 2019 Illinois field experiment. Three genotypes--wild-type (WT), mutant 1 (mt1), and mutant 2 (mt2)--were treated with three treatments: 2% ethanol spray once a day (-1), twice a day (-2), and no treatment (−0, dark green color). B. 2019 Puerto Rico field experiment. Three genotypes—WT (green), mt1(light green), and mt2 (yellow)--were treated with 2% ethanol spray twice a day. C. 2020 Illinois field experiment has two genotypes, WT and mt2, which were treated with 2% ethanol spray twice a day, in addition to mt2 without ethanol treatment (mt2-0, orange). Some genotypes (mt3-mt6), targeting different genes, were not included in this study. Field experiments were set up as a randomized block design. Each plant line was randomized into a single plot based on a random number generator.

**Supplemental Figure 8.**
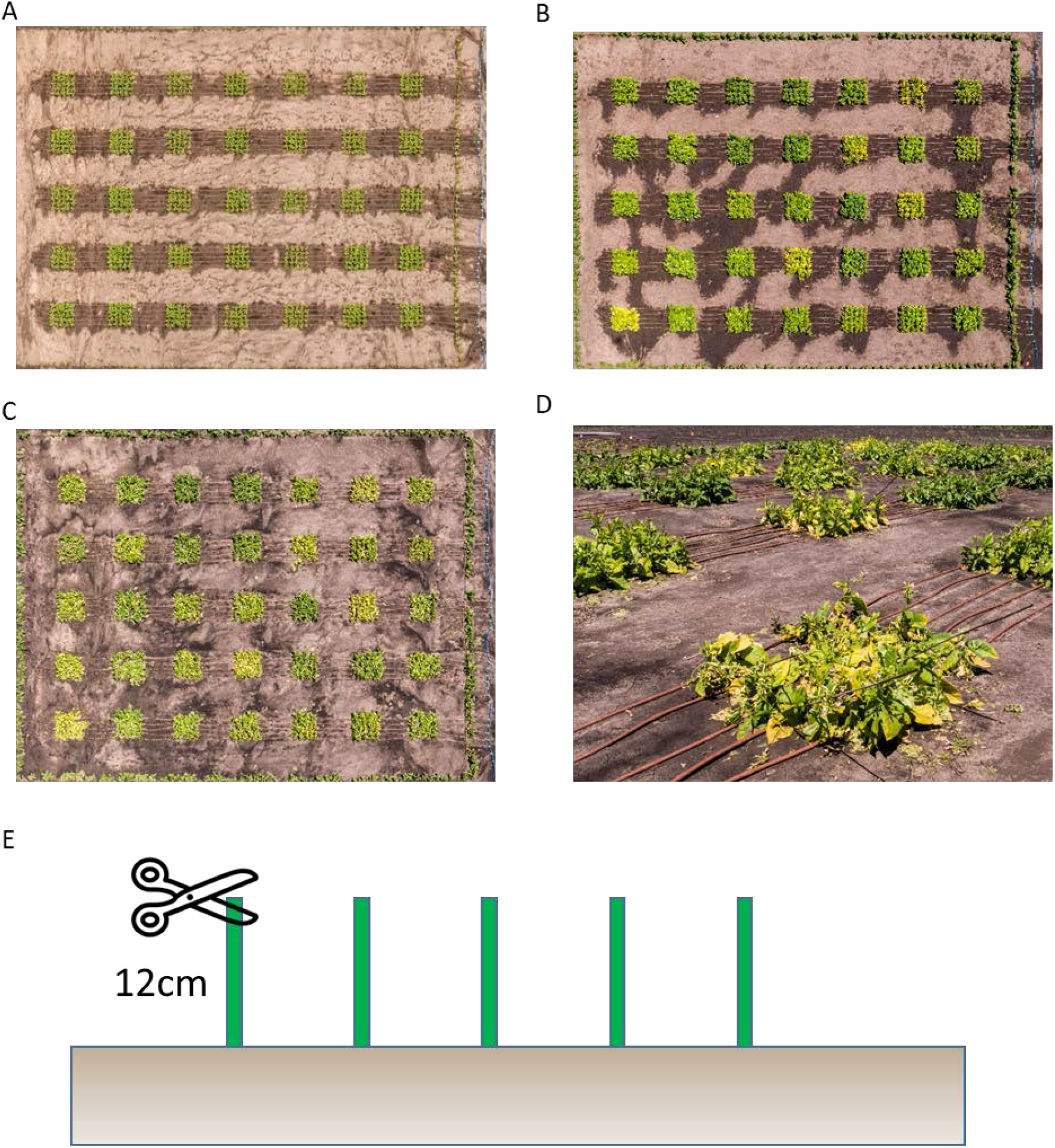
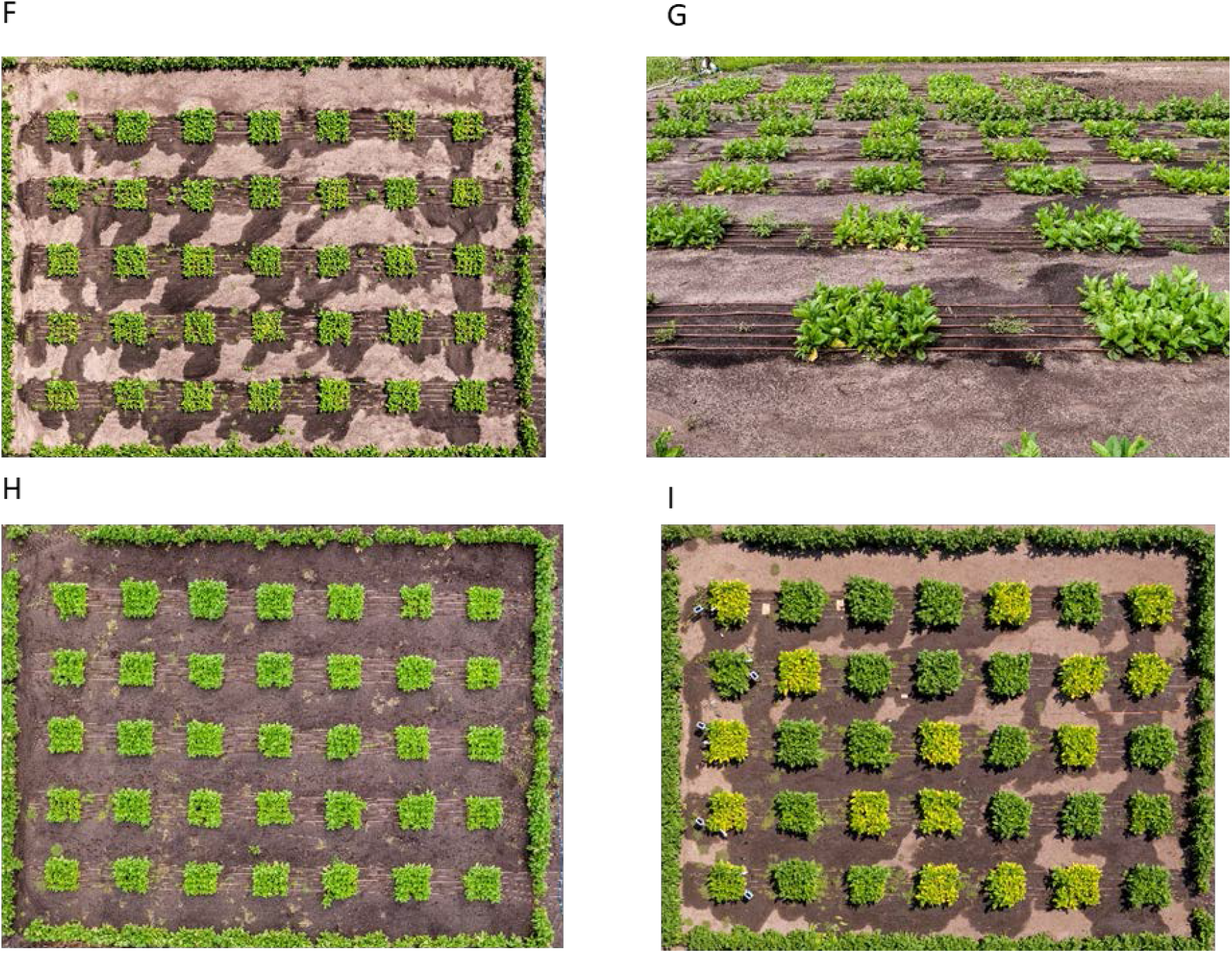
Management of 2020 Illinois field experiment. Aerial pictures of the plot before and after the hail on 11 July 2020. A. Taken on 23 June 2020. B. 10 July 2020. C-D. 13 July 2020. E. Schematic representation of coppice on 14 Jul 2020. All plants were coppiced 12 cm above the ground and allowed let to grow again. F-G. 25 July 2020. H. 31 July 2020. I. 14 August 2020.

**Supplemental Table 1.** Level of total chlorophyll in three field trials. Average total chlorophyll (µg/cm^2^) for each line and treatment shown for each day of measurement. See Dataset 1 for individual measurements. Percentage columns show the reduction in chlorophyll of mt2 compared to the wild-type, e.g., in the IL2020 field trial at 95 DAP (days after planting), mt2-2 showed a 61.49% reduction in chlorophyll compared to WT-2. Ethanol spraying began at 48 DAP (IL2019), 55 DAP (PR2019), and 72 DAP (IL2020) based on timing of canopy closure.

**IL2019**
| Total Chlorophyll Ave |  |  |  |  |  |  |  |  |  |  |  |  |
| --- | --- | --- | --- | --- | --- | --- | --- | --- | --- | --- | --- | --- |
| DAP | mt1-0 | mt1-1 | mt1-2 | mt2-0 | mt2-1 | mt2-2 | WT-0 | WT-1 | WT-2 | mt2-0 vs WT-0 | mt2-1 vs WT-1 | mt2-2 vs WT-2 |
| 38 | 21 | 21.7 | 19.9 | 21.4 | 20 | 21.3 | 20.8 | 23.6 | 22.7 | -2.88% | 15.25% | 6.17% |
| 42 | 25.5 | 26.8 | 26.3 | 27.3 | 26.2 | 27.3 | 29.5 | 26 | 27.5 | 7.46% | -0.77% | 0.73% |
| 46 | 23.6 | 23.2 | 22.4 | 22.9 | 22.2 | 21.8 | 26.3 | 23.8 | 25.4 | 12.93% | 6.72% | 14.17% |
| 48 | 22.2 | 21.9 | 22.5 | 21.2 | 21.3 | 20.7 | 24.1 | 22.4 | 22.9 | 12.03% | 4.91% | 9.61% |
| 52 | 24.5 | 25.1 | 25.7 | 21.4 | 21 | 19.3 | 26.6 | 26.2 | 26.4 | 19.55% | 19.85% | 26.89% |
| 54 | 25.3 | 24.6 | 25.2 | 20.7 | 18.1 | 18.2 | 25.3 | 26.7 | 25.1 | 18.18% | 32.21% | 27.49% |
| 56 | 24.9 | 24.4 | 24 | 20.2 | 14.3 | 14.8 | 27.3 | 27.6 | 26.7 | 26.01% | 48.19% | 44.57% |
| 62 | 25.4 | 24.5 | 25 | 23.8 | 17.7 | 15.9 | 26.3 | 26.2 | 26.6 | 9.51% | 32.44% | 40.23% |
| 67 | 25.3 | 25.6 | 25.6 | 25.3 | 16.3 | 18.1 | 26.6 | 27.1 | 26.3 | 4.89% | 39.85% | 31.18% |
| 69 | 25.2 | 25.3 | 25.5 | 24.4 | 17 | 19.3 | 26.6 | 27.1 | 26 | 8.27% | 37.27% | 25.77% |
| 73 | 25.4 | 24.8 | 25.6 | 24.4 | 19.5 | 20 | 26.3 | 27 | 26.3 | 7.22% | 27.78% | 23.95% |
| 77 | 26.2 | 24.8 | 25.5 | 25.3 | 19.7 | 19.2 | 26.2 | 27.1 | 25.1 | 3.44% | 27.31% | 23.51% |
| 80 | 23.2 | 24.3 | 22.2 | 24.1 | 18.9 | 19.5 | 25.3 | 25.1 | 23.6 | 4.74% | 24.70% | 17.37% |

**PR2019**
| Total Chlorophyll Ave |  |  |  |  |
| --- | --- | --- | --- | --- |
| DAP | mt1 | mt2 | WT | mt2 vs WT |
| 52 | 22 | 23.4 | 24.1 | 2.90% |
| 54 | 19.7 | 22.2 | 22 | -0.91% |
| 56 | 26.2 | 27.2 | 28.5 | 4.56% |
| 66 | 24.6 | 11.9 | 28.5 | 58.25% |

**IL2020**
| Total Chlorophyll Ave |  |  |  |  |
| --- | --- | --- | --- | --- |
| DAP | mt2-0 | mt2-2 | WT-2 | mt2-2 vs WT-2 |
| 72 | 18.53334 | 20.29698 | 21.69 | 6.42% |
| 93 | 27.48573 | 12.05069 | 30.35804 | 60.30% |
| 95 | 27.66678 | 11.69178 | 30.3591 | 61.49% |
| 107 | 30.9555 | 17.35439 | 33.30383 | 47.89% |

**Supplemental Table 2.**
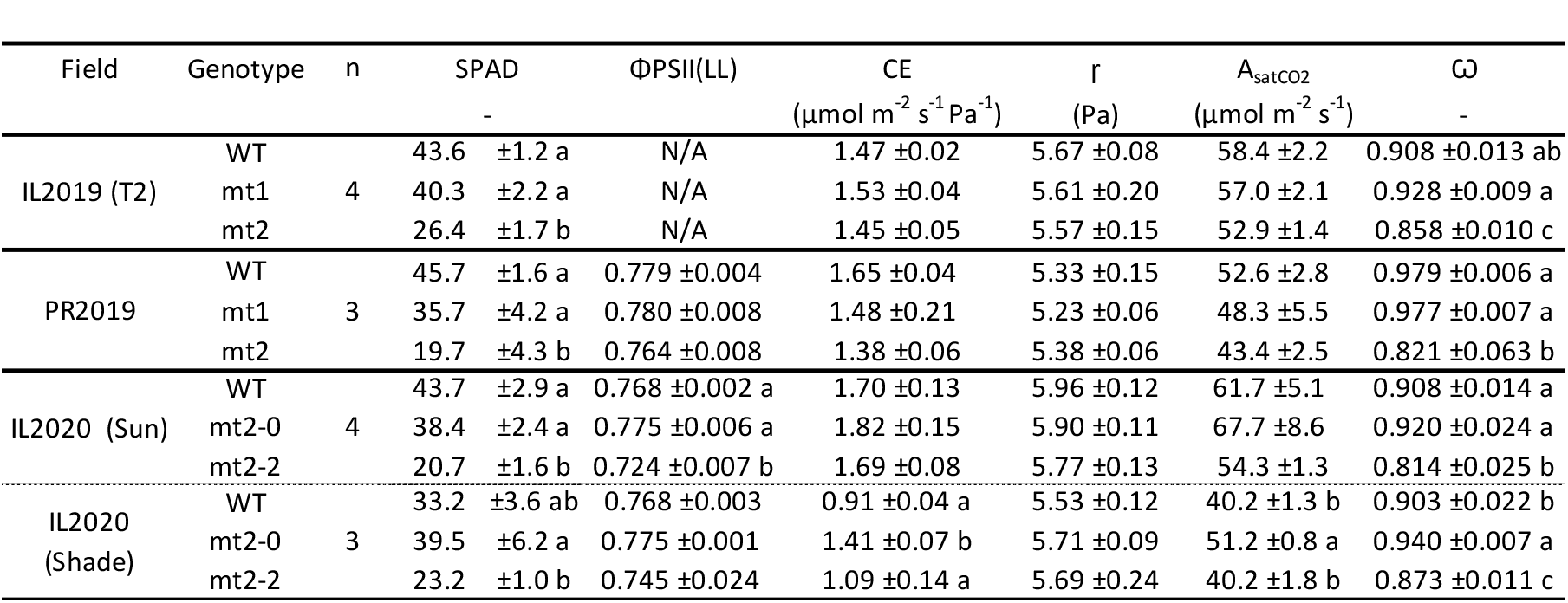
Low and high chlorophyll leaf pigment and photosynthetic parameters from three field trials. The maximum carboxylation efficiency (CE), the CO2 compensation point (Γ), the CO2 saturated rate of Anet (Asat), and the empirical curvature factor for the A/Ci curves (ω) were calculated from A/Ci measurements. Means fitting in linear mixed model are shown with ± SE. Different letters represent significant differences (p < 0.05). N/A represents the data did not fit in the model. *IL 2019 measurements compared genotypes within the twice of a day treatment.

**Supplemental Table 3.** Copy number analysis of plant lines used in this study. Copies_bar represent the copy number of the BASTA gene as detected by quantitative polymerase chain reaction (qRT-PCR) analysis (iDNA Genetics, Norwich, UK). 945-2-27-17 (mt1) and 945-7-11-7 (mt2) are used in the experiments.

| T1 plants |  |  |  | T2 plants |  |  |  |
| --- | --- | --- | --- | --- | --- | --- | --- |
| Sample | Copies_bar | Sample | Copies_bar | Sample | Copies_bar | Sample | Copies_bar |
| 945-2-10 | 1 | 945-7-01 | 6 | 945-2-27-1 | 2 | 945-7-11-13 | 2 |
| 945-2-11 | 2 | 945-7-07 | 6 | 945-2-27-10 | 2 | 945-7-11-1 | 1 |
| 945-2-13 | 1 | 945-7-17 | 6 | 945-2-27-11 | 2 | 945-7-11-10 | 1 |
| 945-2-14 | 0 | 945-7-12 | 5 to 6 | 945-2-27-12 | 2 | 945-7-11-11 | 2 |
| 945-2-15 | 1 | 945-7-10 | 5 to 6 | 945-2-27-13 | 2 | 945-7-11-12 | 1 |
| 945-2-17 | 1 | 945-7-08 | 4 | 945-2-27-14 | 2 | 945-7-11-14 | 1 |
| 945-2-18 | 1 | 945-7-15 | 4 | 945-2-27-15 | 2 | 945-7-11-15 | 2 |
| 945-2-19 | 1 | 945-7-02 | 4 | 945-2-27-16 | 2 | 945-7-11-16 | 2 |
| 945-2-22 | 1 | 945-7-03 | 4 | 945-2-27-17 | No_amplification | 945-7-11-17 | 2 |
| 945-2-23 | 1 | 945-7-18 | 3 | 945-2-27-18 | 2 | 945-7-11-18 | 2 |
| 945-2-25 | 2 | 945-7-09 | 3 | 945-2-27-19 | 2 | 945-7-11-3 | 1 |
| 945-2-26 | 1 | 945-7-05 | 2 | 945-2-27-2 | 2 | 945-7-11-4 | 1 |
| 945-2-27 | 2 | 945-7-14 | 2 | 945-2-27-20 | 2 | 945-7-11-5 | 2 |
| 945-2-28 | 1 | 945-7-13 | 2 | 945-2-27-21 | 2 | 945-7-11-6 | 1 |
| 945-2-29 | 1 | 945-7-11 | 1 | 945-2-27-22 | 2 | 945-7-11-7 | 2 |
| 945-2-30 | 1 | 945-7-16 | 1 | 945-2-27-23 | 2 | 945-7-11-8 | 1 |
| 945-2-31 | 1 |  |  | 945-2-27-24 | 2 | 945-7-11-9 | 2 |
| 945-2-32 | 2 |  |  | 945-2-27-3 | 2 |  |  |
| 945-2-5 | <1 |  |  | 945-2-27-4 | 2 |  |  |
| 945-2-6 | 0 |  |  | 945-2-27-5 | 2 |  |  |
| 945-2-7 | 1 |  |  | 945-2-27-6 | 2 |  |  |
| 945-2-8 | 1 |  |  | 945-2-27-7 | 2 |  |  |
|  |  |  |  | 945-2-27-8 | 2 |  |  |
|  |  |  |  | 945-2-27-9 | 2 |  |  |

**Supplemental Table 4.**
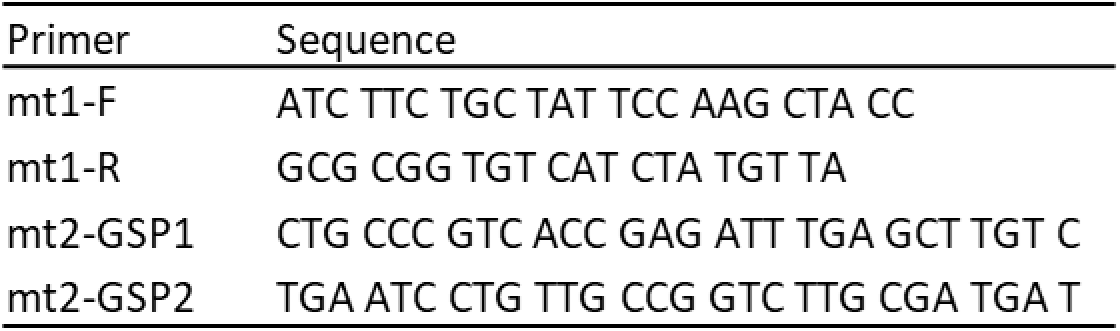
Sequence of primers used in this study.

## Notes

### Competing Interest Statement

The authors have declared no competing interest.

## REFERENCES

Zhu, X. G., Long, S. P., & Ort, D. R. (2010). Improving photosynthetic efficiency for greater yield. Annual Review of Plant Biology, 61, 235–261.

Long, S. P., & Ort, D. R. (2010). More than taking the heat: crops and global change. Current Opinion in Plant Biology, 13(3), 240–247.

Ray, D. K., Mueller, N. D., West, P. C., & Foley, J. A. (2013). Yield trends are insufficient to double global crop production by 2050. PloS One, 8(6), e66428.

Ainsworth, E. A., & Long, S. P. (2021). 30 years of free-air carbon dioxide enrichment (FACE): What have we learned about future crop productivity and its potential for adaptation?. Global Change Biology, 27(1), 27–49.

Ort, D. R., Merchant, S. S., Alric, J., Barkan, A., Blankenship, R. E., Bock, R., … & Zhu, X. G. (2015). Redesigning photosynthesis to sustainably meet global food and bioenergy demand. Proceedings of the National Academy of Sciences, 112(28), 8529–8536.

Simkin, A. J., López-Calcagno, P. E., & Raines, C. A. (2019). Feeding the world: improving photosynthetic efficiency for sustainable crop production. Journal of Experimental Botany, 70(4), 1119–1140.

Sinclair, T. R., & Muchow, R. C. (1999). Radiation use efficiency. Advances in Agronomy, 65, 215–265.

Slattery, R. A., Ainsworth, E. A., & Ort, D. R. (2013). A meta-analysis of responses of canopy photosynthetic conversion efficiency to environmental factors reveals major causes of yield gap. Journal of Experimental Botany, 64(12), 3723–3733.

Slattery, R. A., & Ort, D. R. (2015). Photosynthetic energy conversion efficiency: setting a baseline for gauging future improvements in important food and biofuel crops. Plant Physiology, 168(2), 383–392.

Zhang, D. Y., Sun, G. J., & Jiang, X. H. (1999). Donald’s ideotype and growth redundancy: a game theoretical analysis. Field Crops Research, 61(2), 179–187.

Loomis, R. S. (1993). Optimization theory and crop improvement. International Crop Science I, 583-588.

Denison, R. F., Kiers, E. T., & West, S. A. (2003). Darwinian agriculture: when can humans find solutions beyond the reach of natural selection? The Quarterly Review of Biology, 78(2), 145–168.

Long, S. P. (1993). The significance of light-limiting photosynthesis to crop canopy carbon gain and productivity—a theoretical analysis. In Photosynthesis: photoreactions to plant productivity (pp. 547–559). Springer, Dordrecht.

Long, S. P., Zhu, X. G., Naidu, S. L., & Ort, D. R. (2006). Can improvement in photosynthesis increase crop yields?. Plant, Cell & Environment, 29(3), 315–330.

Ort, D. R., Zhu, X., & Melis, A. (2011). Optimizing antenna size to maximize photosynthetic efficiency. Plant Physiology, 155(1), 79–85.

Walker, B. J., Drewry, D. T., Slattery, R. A., VanLoocke, A., Cho, Y. B., & Ort, D. R. (2018). Chlorophyll can be reduced in crop canopies with little penalty to photosynthesis. Plant Physiology, 176(2), 1215–1232.

Song, Q., Wang, Y., Qu, M., Ort, D. R., & Zhu, X. G. (2017). The impact of modifying photosystem antenna size on canopy photosynthetic efficiency—Development of a new canopy photosynthesis model scaling from metabolism to canopy level processes. Plant, Cell & Environment, 40(12), 2946–2957.

Li, Y., Ren, B., Gao, L., Ding, L., Jiang, D., Xu, X., … & Guo, S. (2013). Less chlorophyll does not necessarily restrain light capture ability and photosynthesis in a chlorophyll-deficient rice mutant. Journal of Agronomy and Crop Science, 199(1), 49–56.

Gu, J., Zhou, Z., Li, Z., Chen, Y., Wang, Z., & Zhang, H. (2017). Rice (*Oryza sativa* L.) with reduced chlorophyll content exhibit higher photosynthetic rate and efficiency, improved canopy light distribution, and greater yields than normally pigmented plants. Field Crops Research, 200, 58–70.

Sakowska, K., Alberti, G., Genesio, L., Peressotti, A., Delle Vedove, G., Gianelle, D., … & Miglietta, F. (2018). Leaf and canopy photosynthesis of a chlorophyll deficient soybean mutant. Plant, Cell & Environment, 41(6), 1427–1437.

Kirst, H., Gabilly, S. T., Niyogi, K. K., Lemaux, P. G., & Melis, A. (2017). Photosynthetic antenna engineering to improve crop yields. Planta, 245(5), 1009–1020.

Felenbok, B., Sequeval, D., Mathieu, M., Sibley, S., Gwynne, D. I., & Davies, R. W. (1988). The ethanol regulon in *Aspergillus nidulans*: characterization and sequence of the positive regulatory gene alcR. Gene, 73(2), 385–396.

Caddick, M. X., Greenland, A. J., Krause, K. P., Qu, N., Riddell, K. V., Salter, M. G., … & Tomsett, A. B. (1998). An ethanol inducible gene switch for plants used to manipulate carbon metabolism. Nature Biotechnology, 16(2), 177.

Salter, M. G., Paine, J. A., Riddell, K. V., Jepson, I., Greenland, A. J., Caddick, M. X., & Tomsett, A. B. (1998). Characterization of the ethanol-inducible alc gene expression system for transgenic plants. The Plant Journal, 16(1), 127–132.

Chen, S., Hofius, D., Sonnewald, U., & Börnke, F. (2003). Temporal and spatial control of gene silencing in transgenic plants by inducible expression of double-stranded RNA. The Plant Journal, 36(5), 731–740.

Good, A. G., Shrawat, A. K., & Muench, D. G. (2004). Can less yield more? Is reducing nutrient input into the environment compatible with maintaining crop production?. Trends in plant science, 9(12), 597–605.

Simmonds, N. W. (1995). The relation between yield and protein in cereal grain. Journal of the Science of Food and Agriculture, 67(3), 309–315.

Feil, B., Thiraporn, R., Getsler, G., & Stamp, P. (1990). Root traits of maize seedlings—indicators of nitrogen efficiency?. In Genetic Aspects of Plant Mineral Nutrition (pp. 97–101). Springer, Dordrecht.

Canevara, M. G., Romani, M., Corbellini, M., Perenzin, M., & Borghi, B. (1994). Evolutionary trends in morphological, physiological, agronomical and qualitative traits of Triticum aestivum L. cultivars bred in Italy since 1900. European Journal of Agronomy, 3(3), 175–185.

Brennan, R. F., Mason, M. G., & Walton, G. H. (2000). Effect of nitrogen fertilizer on the concentrations of oil and protein in canola (Brassica napus) seed. Journal of Plant Nutrition, 23(3), 339–348.

Acreche, M. M., & Slafer, G. A. (2009). Variation of grain nitrogen content in relation with grain yield in old and modern Spanish wheats grown under a wide range of agronomic conditions in a Mediterranean region. The Journal of Agricultural Science, 147(6), 657–667.

Munier-Jolain, N. G., & Salon, C. (2005). Are the carbon costs of seed production related to the quantitative and qualitative performance? An appraisal for legumes and other crops. Plant, Cell & Environment, 28(11), 1388–1395.

Havé, M., Marmagne, A., Chardon, F., & Masclaux-Daubresse, C. (2017). Nitrogen remobilization during leaf senescence: lessons from Arabidopsis to crops. Journal of Experimental Botany, 68(10), 2513–2529

Masclaux-Daubresse, C., Reisdorf-Cren, M., & Orsel, M. (2008). Leaf nitrogen remobilisation for plant development and grain filling. Plant Biology, 10, 23–36.

Uauy, C., Distelfeld, A., Fahima, T., Blechl, A., & Dubcovsky, J. (2006). A NAC gene regulating senescence improves grain protein, zinc, and iron content in wheat. Science, 314(5803), 1298–1301.

Oury, F. X., & Godin, C. (2007). Yield and grain protein concentration in bread wheat: how to use the negative relationship between the two characters to identify favourable genotypes?. Euphytica, 157(1), 45–57.

Gregersen, P. L., Culetic, A., Boschian, L., & Krupinska, K. (2013). Plant senescence and crop productivity. Plant molecular biology, 82(6), 603–622.

Distelfeld, A., Avni, R., & Fischer, A. M. (2014). Senescence, nutrient remobilization, and yield in wheat and barley. Journal of experimental botany, 65(14), 3783–3798.

Campbell, B. W., Mani, D., Curtin, S. J., Slattery, R. A., Michno, J. M., Ort, D. R., … & Stupar, R. M. (2015). Identical substitutions in magnesium chelatase paralogs result in chlorophyll-deficient soybean mutants. G3: Genes, Genomes, Genetics, 5(1), 123-131.

Slattery, R. A., VanLoocke, A., Bernacchi, C. J., Zhu, X. G., & Ort, D. R. (2017). Photosynthesis, light use efficiency, and yield of reduced-chlorophyll soybean mutants in field conditions. Frontiers in Plant Science, 8, 549.

Evans, J. R., & Clarke, V. C. (2019). The nitrogen cost of photosynthesis. Journal of Experimental Botany, 70(1), 7–15.

Kizis, D., Lumbreras, V., & Pagès, M. (2001). Role of AP2/EREBP transcription factors in gene regulation during abiotic stress. FEBS Letters, 498(2-3), 187–189.

Selvaraj, M. G., Jan, A., Ishizaki, T., Valencia, M., Dedicova, B., Maruyama, K., … & Ishitani, M. (2020). Expression of the CCCH-tandem zinc finger protein gene OsTZF5 under a stress-inducible promoter mitigates the effect of drought stress on rice grain yield under field conditions. Plant Biotechnology Journal, 18(8), 1711–1721.

Chen, Y. S., Lo, S. F., Sun, P. K., Lu, C. A., Ho, T. H. D., & Yu, S. M. (2015). A late embryogenesis abundant protein HVA 1 regulated by an inducible promoter enhances root growth and abiotic stress tolerance in rice without yield penalty. Plant Biotechnology Journal, 13(1), 105–116.

Xu, G., Fan, X., & Miller, A. J. (2012). Plant nitrogen assimilation and use efficiency. Annual review of plant biology, 63, 153–182.

Masclaux-Daubresse, C., & Chardon, F. (2011). Exploring nitrogen remobilization for seed filling using natural variation in Arabidopsis thaliana. Journal of Experimental Botany, 62(6), 2131–2142.

Du, S. Y., Zhang, X. F., Lu, Z., Xin, Q., Wu, Z., Jiang, T., … & Zhang, D. P. (2012). Roles of the different components of magnesium chelatase in abscisic acid signal transduction. Plant Molecular Biology, 80(4-5), 519–537.

Tomiyama, M., Inoue, S. I., Tsuzuki, T., Soda, M., Morimoto, S., Okigaki, Y., … & Kinoshita, T. (2014). Mg-chelatase I subunit 1 and Mg-protoporphyrin IX methyltransferase affect the stomatal aperture in Arabidopsis thaliana. Journal of Plant Research, 127(4), 553–563.

Roslan, H. A., Salter, M. G., Wood, C. D., White, M. R., Croft, K. P., Robson, F., … & Caddick, M. X. (2001). Characterization of the ethanol-inducible alc gene-expression system in Arabidopsis thaliana. The Plant Journal, 28(2), 225–235.

Sweetman, J. P., Chu, C., Qu, N., Greenland, A. J., Sonnewald, U., & Jepson, I. (2002). Ethanol vapor is an efficient inducer of the alc gene expression system in model and crop plant species. Plant Physiology, 129(3), 943–948.

Garoosi, G. A., Salter, M. G., Caddick, M. X., & Tomsett, A. B. (2005). Characterization of the ethanol-inducible alc gene expression system in tomato. Journal of Experimental Botany, 56(416), 1635–1642.

Schaarschmidt, S., Qu, N., Strack, D., Sonnewald, U., & Hause, B. (2004). Local induction of the alc gene switch in transgenic tobacco plants by acetaldehyde. Plant and Cell Physiology, 45(11), 1566–1577.

Tomsett, B., Tregova, A., & Caddick, M. (2006). Applications of inducible transcription in plant research and biotechnology. Annual Plant Reviews online, 309-328.

Drew, M. C. (1997). Oxygen deficiency and root metabolism: injury and acclimation under hypoxia and anoxia. Annual Review of Plant Physiology, 48(1), 223–250.

MacDonald, R. C., & Kimmerer, T. W. (1991). Ethanol in the stems of trees. Physiologia Plantarum, 82(4), 582–588.

Junker, B., Wuttke, R., Tiessen, A., Geigenberger, P., Sonnewald, U., & Willmitzer, L. (2004). Temporally regulated expression of a yeast invertase in potato tubers allows dissection of the complex metabolic phenotype obtained following its constitutive expression. Plant Molecular Biology, 56(1), 91–110.

Pesis, E. (2005). The role of the anaerobic metabolites, acetaldehyde and ethanol, in fruit ripening, enhancement of fruit quality and fruit deterioration. Postharvest Biology and Technology, 37(1), 1–19.

Caputo, C., Fatta, N., & Barneix, A. J. (2001). The export of amino acid in the phloem is altered in wheat plants lacking the short arm of chromosome 7B. Journal of Experimental Botany, 52(362), 1761–1768.

Taylor, L., Nunes-Nesi, A., Parsley, K., Leiss, A., Leach, G., Coates, S., … & Hibberd, J. M. (2010). Cytosolic pyruvate, orthophosphate dikinase functions in nitrogen remobilization during leaf senescence and limits individual seed growth and nitrogen content. The Plant Journal, 62(4), 641–652.

Zhao, D., Derkx, A. P., Liu, D. C., Buchner, P., & Hawkesford, M. J. (2015). Overexpression of a NAC transcription factor delays leaf senescence and increases grain nitrogen concentration in wheat. Plant Biology, 17(4), 904–913.

Zhang, L., Tan, Q., Lee, R., Trethewy, A., Lee, Y. H., & Tegeder, M. (2010). Altered xylem-phloem transfer of amino acids affects metabolism and leads to increased seed yield and oil content in Arabidopsis. The Plant Cell, 22(11), 3603–3620.

Srinivasan, V., Kumar, P., & Long, S. P. (2017). Decreasing, not increasing, leaf area will raise crop yields under global atmospheric change. Global Change Biology, 23(4), 1626–1635.

Bogard, M., Allard, V., Brancourt-Hulmel, M., Heumez, E., Machet, J. M., Jeuffroy, M. H., … & Le Gouis, J. (2010). Deviation from the grain protein concentration–grain yield negative relationship is highly correlated to post-anthesis N uptake in winter wheat. Journal of experimental botany, 61(15), 4303–4312.

Palta, J. A., & Fillery, I. R. P. (1995). N application increases pre-anthesis contribution of dry matter to grain yield in wheat grown on a duplex soil. Australian Journal of Agricultural Research, 46(3), 507–518.

Rajcan, I., & Tollenaar, M. (1999). Source: sink ratio and leaf senescence in maize:: II. Nitrogen metabolism during grain filling. Field Crops Research, 60(3), 255–265.

Tegeder, M. (2014). Transporters involved in source to sink partitioning of amino acids and ureides: opportunities for crop improvement. Journal of Experimental Botany, 65(7), 1865–1878.

Myers, S. S., Zanobetti, A., Kloog, I., Huybers, P., Leakey, A. D., Bloom, A. J., … & Usui, Y. (2014). Increasing CO_2_ threatens human nutrition. Nature, 510(7503), 139–142.

Loladze, I. (2002). Rising atmospheric CO2 and human nutrition: toward globally imbalanced plant stoichiometry? Trends in Ecology & Evolution, 17(10), 457–461.

Mcgrath, J. M., & Lobell, D. B. (2013). Reduction of transpiration and altered nutrient allocation contribute to nutrient decline of crops grown in elevated CO2 concentrations. Plant, Cell & Environment, 36(3), 697–705.

Engler, C., Kandzia, R., & Marillonnet, S. (2008). A one pot, one step, precision cloning method with high throughput capability. PloS One, 3(11), e3647.

Engler, C., Gruetzner, R., Kandzia, R., & Marillonnet, S. (2009). Golden gate shuffling: a one-pot DNA shuffling method based on type IIs restriction enzymes. PloS One, 4(5), e5553.

Gallois, P., & Marinho, P. (1995). Leaf disk transformation using *Agrobacterium tumefaciens*-expression of heterologous genes in tobacco. Methods in Molecular Biology, 49, 39–48.

Cho, Y. B., Jones, S. I., & Vodkin, L. O. (2019). Nonallelic homologous recombination events responsible for copy number variation within an RNA silencing locus. Plant Direct, 3(8), e00162.

De Vree, P. J., De Wit, E., Yilmaz, M., Van De Heijning, M., Klous, P., Verstegen, M. J., … & De Laat, W. (2014). Targeted sequencing by proximity ligation for comprehensive variant detection and local haplotyping. Nature Biotechnology, 32(10), 1019–1025.

South, P. F., Cavanagh, A. P., Liu, H. W., & Ort, D. R. (2019). Synthetic glycolate metabolism pathways stimulate crop growth and productivity in the field. Science, 363(6422).

Kromdijk, J., Głowacka, K., Leonelli, L., Gabilly, S. T., Iwai, M., Niyogi, K. K., & Long, S. P. (2016). Improving photosynthesis and crop productivity by accelerating recovery from photoprotection. Science, 354(6314), 857–861.

Ritchie, R. J. (2006). Consistent sets of spectrophotometric chlorophyll equations for acetone, methanol and ethanol solvents. Photosynthesis Research, 89(1), 27–41.

Wang, C. S., & Vodkin, L. O. (1994). Extraction of RNA from tissues containing high levels of procyanidins that bind RNA. Plant Molecular Biology Reporter, 12(2), 132–145.

Cho, Y. B., Jones, S. I., & Vodkin, L. (2013). The transition from primary siRNAs to amplified secondary siRNAs that regulate chalcone synthase during development of *Glycine max* seed coats. PLoS One, 8(10), e76954.

Cho, Y. B., Jones, S. I., & Vodkin, L. O. (2017). Mutations in Argonaute5 illuminate epistatic interactions of the K1 and I loci leading to saddle seed color patterns in Glycine max. The Plant Cell, 29(4), 708–725.

Edwards, K. D., Fernandez-Pozo, N., Drake-Stowe, K., Humphry, M., Evans, A. D., Bombarely, A., … & Mueller, L. A. (2017). A reference genome for *Nicotiana tabacum* enables map-based cloning of homeologous loci implicated in nitrogen utilization efficiency. BMC Genomics, 18(1), 1–14.

Patro, R., Duggal, G., Love, M. I., Irizarry, R. A., & Kingsford, C. (2017). Salmon provides fast and bias-aware quantification of transcript expression. Nature methods, 14(4), 417–419.

Masters, M. D., Black, C. K., Kantola, I. B., Woli, K. P., Voigt, T., David, M. B., & DeLucia, E. H. (2016). Soil nutrient removal by four potential bioenergy crops: Zea mays, Panicum virgatum, Miscanthus× giganteus, and prairie. Agriculture, Ecosystems & Environment, 216, 51–60.

