## Supplementary material for "Reducing chlorophyll level in seed filling stages results in higher seed nitrogen without impacting canopy carbon assimilation": SI appendix: Fig1.R

```
#2018 greenhouse
#reference

#packages that need
library(ggplot2)
library(ggpubr)
library(dplyr)

#read data
chl <-read.csv("greenhouse_test.csv")

#ggplot with jitter
ggboxplot(chl, x = "DAS", y = "Total.Chlorophyll", color = "Treatment", size = 1.2,
          ylab="Total Chlorophyll (�g/cm2)", xlab = "Day after application", add = "jitter", 
          palette = "RuBp")
```
