## Supplementary material for "Reducing chlorophyll level in seed filling stages results in higher seed nitrogen without impacting canopy carbon assimilation": SI appendix: Fig1_2.R

```
#2019 chlorophyll sRNA and CHL level
#reference

#packages that need
library(ggplot2)
library(ggpubr)
library(dplyr)
library(car)

#read data
sRNA_il2019 <-read.delim("YC_sRNA_Gene_level_counts_Full_Analysis.txt")

#subset data
library("stringr")  
chelatase <- sRNA_il2019[str_detect(sRNA_il2019$GENENAME, "chelatase"),]
write.csv(chelatase,'chelatase.csv')

#sRNA
sRNA<-read.csv("Nitab4.5_0006237g0030.csv")

ggline(sRNA, x = "Time1", y = "sRNA", ylab="sRNA level (CPM)", color = "sample", xlab = "Hour after spray", add = "mean_se",
       palette = c("#E7B800"), size=1) 
#ratio 300:180

#a/b ratio
abratio <-read.csv("chlrophyll_abratio.csv")

ggline(abratio, x = "Time", y = "abratio", color = "sample", linetype = "sample",
       shape = "sample", ylab="Chlorophyll a/b", xlab = "Hour after spray", 
       add = "mean_se", palette = c("#E7B800","#E7B800","green4"), size=1)

#total chlorophyll
chl <-read.csv("chlrophyll_chl.csv")

ggline(chl, x = "Time", y = "chl", color = "sample", linetype = "sample",
       shape = "sample", ylab="Total chlorophyll (�g/\u33A0)", xlab = "Hour after spray", 
       add = "mean_se", palette = c("#E7B800","#E7B800","green4"), size=1)
#ratio 400:200
```
