## Supplementary material for "Reducing chlorophyll level in seed filling stages results in higher seed nitrogen without impacting canopy carbon assimilation": SI appendix: Fig2B.R

```
#Loading packages
library(ggplot2)
library(ggpubr)
library(dplyr)
library(car)
library(lme4)
library(lmerTest)
library(multcomp)
library(lsmeans)
library(simr)

#read data
Biomass<-read.csv("Fig2B.csv")

#average of plot
Biomass_average<-group_by(Biomass, Treatment, Genotype, Block) %>%
  summarise(Weight=mean(Weight,na.rm=T)
  )

################################################################################################################################
#Complete Randomized block design

################################################################################################################################
#Evaluate Genotype treatment interaction

# 2.Model fitting
Biomass_average.lmer <- lmer(Weight ~ Treatment + Genotype + Genotype*Treatment + (1 | Block), Biomass_average, REML = TRUE)  
anova(Biomass_average.lmer)
summary(Biomass_average.lmer)
ranova(Biomass_average.lmer)

# 3. Model evaluation
#Residuals
plot(Biomass_average.lmer)
qqnorm(resid(Biomass_average.lmer))
qqline(resid(Biomass_average.lmer))
ggqqplot(resid(Biomass_average.lmer))


# 4. Parameter estimates
summary(Biomass_average.lmer)
confint(Biomass_average.lmer)
anova(Biomass_average.lmer, type=3)
anova(Biomass_average.lmer, type=3, ddf = "Kenward-Roger") 

#We failed to find treatment, genotype or interaction effect on above-ground biomass.

###########################
##Evaluate Treatment effect within WT
Biomass<-read.csv("Fig2B.csv")
Biomass<-subset(Biomass, Genotype=="wt")

#average
Biomass_average<-group_by(Biomass, Treatment, Block) %>%
  summarise(Weight=mean(Weight,na.rm=T)
  )
#ggplot with jitter
ggboxplot(Biomass_average, x = "Treatment", y = "Weight", color = "Treatment", size = 1.2,
          ylab="Value", xlab = "Treatment", add = "jitter", 
          palette = c("limegreen","#E7B800","green4"))

# 1.Data analysis. 
#Is the data skewed?
boxplot(Weight~ Treatment, Biomass_average)

group_by(Biomass_average, Treatment) %>%
  summarise(
    n = n(),
    mean = mean(Weight, na.rm = TRUE),
    sd = sd(Weight, na.rm = TRUE),
    se = sd / sqrt(n)
  )

#Is this data balanced?
replications(Weight ~ Treatment + Block, data = Biomass_average)
!is.list(replications(Weight ~ Treatment  + Block, data = Biomass_average))
interaction.plot(Biomass_average$Block, Biomass_average$Treatment, Biomass_average$Weight) 


#Is the interaction between random and fixed effect?
with(Biomass_average, interaction.plot(Treatment, Block, Weight)) 
residualPlots(lm(Weight ~ Block + Treatment, Biomass_average)) 


# 2.Model fitting

Biomass_average.lmer <- lmer(Weight ~ Treatment + (1 | Block), Biomass_average, REML = TRUE)  
anova(Biomass_average.lmer)
summary(Biomass_average.lmer)
ranova(Biomass_average.lmer)

# 3. Model evaluation
#Residuals
plot(Biomass_average.lmer)
qqnorm(resid(Biomass_average.lmer))
qqline(resid(Biomass_average.lmer))
ggqqplot(resid(Biomass_average.lmer))


# 4. Parameter estimates
summary(Biomass_average.lmer)
confint(Biomass_average.lmer)
anova(Biomass_average.lmer, type=3)
anova(Biomass_average.lmer, type=3, ddf = "Kenward-Roger") 


# 5. planned comparisons and pairwise post-hoc tests

summary(glht(Biomass_average.lmer, linfct = mcp(Treatment ="Tukey")))

lsmeans(Biomass_average.lmer, pairwise ~ Treatment)
summary(glht(Biomass_average.lmer, linfct = lsm(pairwise ~ Treatment)))
confint(glht(Biomass_average.lmer, linfct = lsm(pairwise ~ Treatment)))


#6. power analysis for linear mixed model (https://www.r-bloggers.com/2017/07/power-analysis-and-sample-size-calculation-for-agriculture/)
library(simr)
summary(Biomass_average.lmer)
powerSim(Biomass_average.lmer, alpha=0.05, nsim=1000)

#calculate sample size (https://rstudio-pubs-static.s3.amazonaws.com/484106_6b51212f20164fdd88cd7cce89bdef79.html)

#Changing the number of classes
model_ext_class <- extend(Biomass_average.lmer, along="Block", n=50)
model_ext_class

#Power curve 
set.seed(0)
p_curve <- powerCurve(model_ext_class, along="Block",breaks=c(10,20,30,40,50),
                      nsim=500,alpha=.05, progress=FALSE)
plot(p_curve)

#We failed to find treatment effect on above-ground biomass of WT. 
#On the other word, we could not find that ethanol spray affect above ground biomass of WT.

###########################
##Evaluate treatment effect in mt2 which showed low chlorophyll level
Biomass<-read.csv("Fig2B.csv")
Biomass<-subset(Biomass, Genotype=="mt2")

#average
Biomass_average<-group_by(Biomass, Treatment, Block) %>%
  summarise(Weight=mean(Weight,na.rm=T)
  )
#ggplot with jitter
ggboxplot(Biomass_average, x = "Treatment", y = "Weight", color = "Treatment", size = 1.2,
          ylab="Value", xlab = "Treatment", add = "jitter", 
          palette = c("limegreen","#E7B800","green4"))

# 1.Data analysis. 
#Is the data skewed?
boxplot(Weight~ Treatment, Biomass_average)

group_by(Biomass_average, Treatment) %>%
  summarise(
    n = n(),
    mean = mean(Weight, na.rm = TRUE),
    sd = sd(Weight, na.rm = TRUE),
    se = sd / sqrt(n)
  )

#Is this data balanced?
replications(Weight ~ Treatment + Block, data = Biomass_average)
!is.list(replications(Weight ~ Treatment  + Block, data = Biomass_average))
interaction.plot(Biomass_average$Block, Biomass_average$Treatment, Biomass_average$Weight) 


#Is the interaction between random and fixed effect?
with(Biomass_average, interaction.plot(Treatment, Block, Weight)) 
residualPlots(lm(Weight ~ Block + Treatment, Biomass_average)) 


# 2.Model fitting

Biomass_average.lmer <- lmer(Weight ~ Treatment + (1 | Block), Biomass_average, REML = TRUE)  
anova(Biomass_average.lmer)
summary(Biomass_average.lmer)
ranova(Biomass_average.lmer)

# 3. Model evaluation
#Residuals
plot(Biomass_average.lmer)
qqnorm(resid(Biomass_average.lmer))
qqline(resid(Biomass_average.lmer))
ggqqplot(resid(Biomass_average.lmer))


# 4. Parameter estimates
summary(Biomass_average.lmer)
confint(Biomass_average.lmer)
anova(Biomass_average.lmer, type=3)
anova(Biomass_average.lmer, type=3, ddf = "Kenward-Roger") 


# 5. planned comparisons and pairwise post-hoc tests

summary(glht(Biomass_average.lmer, linfct = mcp(Treatment ="Tukey")))

lsmeans(Biomass_average.lmer, pairwise ~ Treatment)
summary(glht(Biomass_average.lmer, linfct = lsm(pairwise ~ Treatment)))
confint(glht(Biomass_average.lmer, linfct = lsm(pairwise ~ Treatment)))


#6. power analysis for linear mixed model (https://www.r-bloggers.com/2017/07/power-analysis-and-sample-size-calculation-for-agriculture/)
library(simr)
summary(Biomass_average.lmer)
powerSim(Biomass_average.lmer, alpha=0.05, nsim=1000)

#calculate sample size (https://rstudio-pubs-static.s3.amazonaws.com/484106_6b51212f20164fdd88cd7cce89bdef79.html)

#Changing the number of classes
model_ext_class <- extend(Biomass_average.lmer, along="Block", n=50)
model_ext_class

#Power curve 
set.seed(0)
p_curve <- powerCurve(model_ext_class, along="Block",breaks=c(10,20,30,40,50),
                      nsim=500,alpha=.05, progress=FALSE)
plot(p_curve)

#We failed to find the effect of treatment on above ground biomass of mt2.
#on the other word, we failed to find effect of low chlorophyll (50% reduction) on above-ground biomass of mt2.

###########################
##Evaluate genotype effect within treatment
Biomass<-read.csv("Fig2B.csv")
Biomass<-subset(Biomass, Treatment=="0")

#average
Biomass_average<-group_by(Biomass, Genotype, Block) %>%
  summarise(Weight=mean(Weight,na.rm=T)
  )
#ggplot with jitter
ggboxplot(Biomass_average, x = "Genotype", y = "Weight", color = "Genotype", size = 1.2,
          ylab="Value", xlab = "Genotype", add = "jitter", 
          palette = c("limegreen","#E7B800","green4"))

# 1.Data analysis. 
#Is the data skewed?
boxplot(Weight~ Genotype, Biomass_average)

group_by(Biomass_average, Genotype) %>%
  summarise(
    n = n(),
    mean = mean(Weight, na.rm = TRUE),
    sd = sd(Weight, na.rm = TRUE),
    se = sd / sqrt(n)
  )

#Is this data balanced?
replications(Weight ~ Genotype + Block, data = Biomass_average)
!is.list(replications(Weight ~ Genotype  + Block, data = Biomass_average))
interaction.plot(Biomass_average$Block, Biomass_average$Genotype, Biomass_average$Weight) 


#Is the interaction between random and fixed effect?
with(Biomass_average, interaction.plot(Genotype, Block, Weight)) 
residualPlots(lm(Weight ~ Block + Genotype, Biomass_average)) 


# 2.Model fitting

Biomass_average.lmer <- lmer(Weight ~ Genotype + (1 | Block), Biomass_average, REML = TRUE)  
anova(Biomass_average.lmer)
summary(Biomass_average.lmer)
ranova(Biomass_average.lmer)

# 3. Model evaluation
#Residuals
plot(Biomass_average.lmer)
qqnorm(resid(Biomass_average.lmer))
qqline(resid(Biomass_average.lmer))
ggqqplot(resid(Biomass_average.lmer))


# 4. Parameter estimates
summary(Biomass_average.lmer)
confint(Biomass_average.lmer)
anova(Biomass_average.lmer, type=3)
anova(Biomass_average.lmer, type=3, ddf = "Kenward-Roger") 


# 5. planned comparisons and pairwise post-hoc tests

summary(glht(Biomass_average.lmer, linfct = mcp(Genotype ="Tukey")))

lsmeans(Biomass_average.lmer, pairwise ~ Genotype)
summary(glht(Biomass_average.lmer, linfct = lsm(pairwise ~ Genotype)))
confint(glht(Biomass_average.lmer, linfct = lsm(pairwise ~ Genotype)))


#6. power analysis for linear mixed model (https://www.r-bloggers.com/2017/07/power-analysis-and-sample-size-calculation-for-agriculture/)
library(simr)
summary(Biomass_average.lmer)
powerSim(Biomass_average.lmer, alpha=0.05, nsim=1000)

#calculate sample size (https://rstudio-pubs-static.s3.amazonaws.com/484106_6b51212f20164fdd88cd7cce89bdef79.html)

#Changing the number of classes
model_ext_class <- extend(Biomass_average.lmer, along="Block", n=50)
model_ext_class

#Power curve 
set.seed(0)
p_curve <- powerCurve(model_ext_class, along="Block",breaks=c(10,20,30,40,50),
                      nsim=500,alpha=.05, progress=FALSE)
plot(p_curve)

#We tested genotype effect in three treatments (0, 1, 2). We failed to find genotype effect on above-ground biomass.

#Overall, we failed to find effect of low chlorophyll or ethanol effect on above-ground biomass.
```
