## Supplementary material for "Reducing chlorophyll level in seed filling stages results in higher seed nitrogen without impacting canopy carbon assimilation": SI appendix: Fig2B_ggplot.R

```
#Loading packages
library(ggplot2)
library(ggpubr)
library(dplyr)
library(car)
library(lme4)
library(lmerTest)
library(multcomp)
library(lsmeans)
library(simr)

##########################################################################################################################
#Display data points
#http://www.sthda.com/english/articles/24-ggpubr-publication-ready-plots/76-add-p-values-and-significance-levels-to-ggplots/
#read data
Biomass<-read.csv("Fig2B.csv")

#ggplot with jitter to show every value within blocks
#need to make a genotype-treatment column
ggboxplot(Biomass, x = "Treatment", y = "Weight", color = "Genotype", size = 1.2,
          ylab="Above ground biomass(g)", xlab = "Treatment", add = "jitter", 
          palette = c("limegreen","#E7B800","green4"))

#Prepare the data : convert Block and Genotype to factor variables
Biomass$Block<-as.factor(Biomass$Block)
Biomass$Genotype.Treatment<-as.factor(Biomass$Genotype)

theme_set(
  theme_classic() +
    theme(legend.position = "top")
)
e<-ggplot(Biomass, aes(x=Genotype.Treatment, y=Weight, color=Genotype))

#jitterplot
e + geom_jitter(
  aes(shape = Block, color = Genotype.Treatment), size = 2,
  position = position_jitterdodge(jitter.width = 0.2, dodge.width = 0.8)
) +
  stat_summary(
    aes(color = Genotype.Treatment), fun.data="mean_sdl", fun.args = list(mult=1), 
    size = 0.8, position = position_dodge(0.8)
  )+
  scale_color_manual(values =  c("limegreen","#E7B800","green4","limegreen","#E7B800","green4","limegreen","#E7B800","green4"))+
  labs(y="Dry above-ground biomass (g)", x="Genotype-Treatment")+
  theme (legend.title= element_text(size=10), legend.text= element_text(size=10), 
         axis.title.x = element_text(size=15), axis.title.y = element_text(size=15))
```
