## Supplementary material for "Reducing chlorophyll level in seed filling stages results in higher seed nitrogen without impacting canopy carbon assimilation": SI appendix: Fig2C.R

```
#2019 IL chlorophyll
#reference

#packages that need
library(ggplot2)
library(ggpubr)
library(dplyr)
library(car)
library(lme4)
library(lsmeans)

#read data for correlation between spad and chlorophyll (ug/cm^2)
chl_spad<-read.csv("chl_spad.csv")

model <- lm(Total.Chlorophyll~SPAD, data = chl_spad)
model
summary (model)

#Total Chlorophyll (µg/cm^2)= 0.6390*SPAD-0.9306

model_a <- lm(Chlorophyll.a~SPAD, data = chl_spad)
model_a
summary(model_a)

#read whole season data

chl_ws_il2019 <-read.csv("IL2019_chl_whole_season.csv")

Biomass$Block<-as.factor(Biomass$Block)
Biomass$Genotype.Treatment<-as.factor(Biomass$Genotype.Treatment)

ggline(chl_ws_il2019, x = "DAP", y = "Total.Chlorophyll", color = "Genotype.Treatment", linetype = "Genotype.Treatment",
       shape = "Genotype.Treatment", ylab="Total Chlorophyll (µg/\u33A0)", xlab = "Days after plant", 
       add = "mean_se", palette = c("limegreen","limegreen","limegreen","#E7B800","#E7B800","#E7B800","green4","green4","green4"), size=1) +
  scale_shape_manual(values = 0:9) +
  theme(legend.position = "right")+
  theme (legend.title= element_text(size=10), legend.text= element_text(size=10), 
         axis.title.x = element_text(size=15), axis.title.y = element_text(size=15))

#subset data
chl_before_spray <- subset(chl_ws_il2019, DAS<4)
chl_after_spray <- subset(chl_ws_il2019, DAS>4)

group_by(chl_before_spray, Genotype.Treatment) %>%
  summarise(
    count = n(),
    mean = mean(Total.Chlorophyll, na.rm = TRUE),
    sd = sd(Total.Chlorophyll, na.rm = TRUE),
    se = sd/sqrt(n)
  )

group_by(chl_after_spray, Genotype.Treatment) %>%
  summarise(
    count = n(),
    mean = mean(Total.Chlorophyll, na.rm = TRUE),
    sd = sd(Total.Chlorophyll, na.rm = TRUE)
  )

#ggplot with jitter
ggboxplot(chl, x = "Treatment", y = "Total.Chlorophyll", color = "Genotype", size = 1.2,
          ylab="Total Chlorophyll (µg/cm2)", xlab = "Treatment", add = "jitter", palette = c("limegreen","gold","green4"))


#Randomized complete block design test. fixed effect: genotype and treatment. block is random effect.
#data were compared by linear mixed model analysis using the lmer function and type III ANOVA to account for block effects 
#using eight plants per genotype for n=4 blocks

#compare models
data.chl.lmer <- lmer(Total.Chlorophyll ~ Genotype*Treatment + (1|Block), data=chl, REML=F) 


Anova(data.chl.lmer, type = 3, white.adjust=TRUE)

#model evaluation looks OK
ggqqplot(resid(data.chl.lmer))

#parameter estimates
summary(data.chl.lmer)
confint(data.chl.lmer)
anova(data.chl.lmer)

#post-hoc test
library(multcomp)
summary(glht(data.chl.lmer, linfct = mcp(Genotype = "Tukey")))
```
