## Supplementary material for "Reducing chlorophyll level in seed filling stages results in higher seed nitrogen without impacting canopy carbon assimilation": SI appendix: Fig2E_v1.R

```
#2019 PR chlorophyll
#reference

#packages that need
library(ggplot2)
library(ggpubr)
library(dplyr)
library(car)
library(lme4)
library(lsmeans)

#Total Chlorophyll (µg/cm^2)= 0.6390*SPAD-0.9306

#read data
chl_pr2019 <-read.csv("PR_chlorophyll.csv")

ggline(chl_pr2019, x = "DAP", y = "Total.Chlorophyll", color = "Genotype", linetype = "Genotype",
       shape = "Genotype", ylab="Total chlorophyll (µg/\u33A0)", xlab = "Day after plant", 
       add = "mean_se", palette = c("limegreen","gold","green4"), font.label = list(size = 1, color = "black"), size=1.2) +
  theme (legend.title= element_text(size=10), legend.text= element_text(size=10), 
         axis.title.x = element_text(size=15), axis.title.y = element_text(size=15))
  

#Randomized complete block design test. fixed effect: genotype and treatment. block is random effect.
#data were compared by linear mixed model analysis using the lmer function and type III ANOVA to account for block effects 
#using eight plants per genotype for n=4 blocks


#ggplot with jitter
ggboxplot(chl_pr2019, x = "DAP", y = "Total.Chlorophyll", color = "Genotype",
          ylab="Total Chlorophyll (µg/cm2)", xlab = "Days after plant", add = "jitter", 
          palette = c("limegreen","#E7B800","green4"), size=1)

#compare models
data.chl.lmer <- lmer(Total.Chlorophyll ~ Genotype + (1|Block), data=chl_10, REML=F) 

Anova(data.chl.lmer, type = 3, white.adjust=TRUE)

#model evaluation looks OK
ggqqplot(resid(data.chl.lmer))

#parameter estimates
summary(data.chl.lmer)
confint(data.chl.lmer)

#post-hoc test
library(multcomp)
summary(glht(data.chl.lmer, linfct = mcp(Genotype = "Tukey")))

#subset data
chl_before_spray <- subset(chl_pr2019, DAP<57)
chl_after_spray <- subset(chl_pr2019, DAP>57)

group_by(chl_before_spray, Genotype) %>%
  summarise(
    count = n(),
    mean = mean(Total.Chlorophyll, na.rm = TRUE),
    sd = sd(Total.Chlorophyll, na.rm = TRUE)
  )

group_by(chl_after_spray, Genotype) %>%
  summarise(
    count = n(),
    mean = mean(Total.Chlorophyll, na.rm = TRUE),
    sd = sd(Total.Chlorophyll, na.rm = TRUE)
  )

data.chl.lmer <- lmer(Total.Chlorophyll ~ Genotype + (1|Block), data=chl_after_spray, REML=F) 

Anova(data.chl.lmer, type = 3, white.adjust=TRUE)
```
