## Supplementary material for "Reducing chlorophyll level in seed filling stages results in higher seed nitrogen without impacting canopy carbon assimilation": SI appendix: Fig2F.R

```
#Loading packages
library(ggplot2)
library(ggpubr)
library(dplyr)
library(car)
library(lme4)
library(lmerTest)
library(multcomp)
library(lsmeans)
library(simr)

#PR2019, we did not look at treatment effect as IL2019 showed no ethanol effect on above-ground biomass.
CN<-read.csv("Fig2F.csv")

#average of plot
CN<-group_by(CN, Genotype, Block) %>%
  summarise(Weight=mean(Weight,na.rm=T)
  )

################################################################################################################################
#Complete Randomized block design

################################################################################################################################
#Weight
#ggplot with jitter
ggboxplot(CN, x = "Genotype", y = "Weight", color = "Genotype", size = 1.2,
          ylab="Value", xlab = "Genotype", add = "jitter", 
          palette = c("limegreen","#E7B800","green4"))

# 1.Data analysis. 
#Is the data skewed?
boxplot(Weight~ Genotype, CN)

group_by(CN, Genotype) %>%
  summarise(
    n = n(),
    mean = mean(Weight, na.rm = TRUE),
    sd = sd(Weight, na.rm = TRUE),
    se = sd / sqrt(n)
  )

#Is this data balanced?
replications(Weight ~ Genotype + Block, data = CN)
!is.list(replications(Weight ~ Genotype + Block, data = CN))
interaction.plot(CN$Block, CN$Genotype, CN$Weight) 


#Is the interaction between random and fixed effect?
with(CN, interaction.plot(Genotype, Block, Weight)) 
residualPlots(lm(Weight ~ Block + Genotype, CN)) 


# 2.Model fitting
CN.lmer <- lmer(Weight ~ Genotype + (1 | Block), CN, REML = TRUE)  
anova(CN.lmer)
summary(CN.lmer)
ranova(CN.lmer)

# 3. Model evaluation
#Residuals
plot(CN.lmer)
qqnorm(resid(CN.lmer))
qqline(resid(CN.lmer))
ggqqplot(resid(CN.lmer))


# 4. Parameter estimates
summary(CN.lmer)
confint(CN.lmer)
anova(CN.lmer, type=3)
anova(CN.lmer, type=3, ddf = "Kenward-Roger") 


# 5. planned comparisons and pairwise post-hoc tests

summary(glht(CN.lmer, linfct = mcp(Genotype ="Tukey")))

lsmeans(CN.lmer, pairwise ~ Genotype)
summary(glht(CN.lmer, linfct = lsm(pairwise ~ Genotype)))
confint(glht(CN.lmer, linfct = lsm(pairwise ~ Genotype)))


#6. power analysis for linear mixed model (https://www.r-bloggers.com/2017/07/power-analysis-and-sample-size-calculation-for-agriculture/)
library(simr)
summary(CN.lmer)
powerSim(CN.lmer, alpha=0.05, nsim=1000)

#calculate sample size (https://rstudio-pubs-static.s3.amazonaws.com/484106_6b51212f20164fdd88cd7cce89bdef79.html)

#Changing the number of classes
model_ext_class <- extend(CN.lmer, along="Block", n=50)
model_ext_class

#Power curve 
set.seed(0)
p_curve <- powerCurve(model_ext_class, along="Block",breaks=c(10,20,30,40,50),
                      nsim=500,alpha=.05, progress=FALSE)
plot(p_curve)

## We failed to find low-chlorophyll effect on above-ground biomass. 
## p-value is lower than one of IL 2019 test. We tested mt2 with treatment and without treatment in 2020 IL. 
```
