## Supplementary material for "Reducing chlorophyll level in seed filling stages results in higher seed nitrogen without impacting canopy carbon assimilation": SI appendix: Fig2F_ggplot.R

```
#2019 PR field trial
#reference

#packages that need
library(ggplot2)
library(ggpubr)
library(dplyr)
library(car)
library(lme4)

#read data
Biomass<-read.csv("PR_Biomass_individual1.csv")

#ggplot with jitter
ggboxplot(Biomass, x = "Genotype", y = "Total", color = "Genotype", size = 1.2,
          ylab="Above ground biomass(g)", xlab = F, add = "jitter", order=c("mt1","mt2","WT"),
          palette = c("limegreen","#E7B800","green4"), short.panel.labs = F)
#Prepare the data : convert Block and Genotype to factor variables
Biomass$Block<-as.factor(Biomass$Block)
Biomass$Genotype<-as.factor(Biomass$Genotype)

theme_set(
  theme_classic() +
    theme(legend.position = "top")
)
e<-ggplot(Biomass, aes(x=Genotype, y=Weight, color=Block))

#jitterplot
e + geom_jitter(
  aes(shape = Block, color = Genotype), size = 2,
  position = position_jitterdodge(jitter.width = 0.2, dodge.width = 0.8)
) +
  stat_summary(
    aes(color = Genotype), fun.data="mean_sdl", fun.args = list(mult=1), 
    size = 0.8, position = position_dodge(0.8)
  )+
  scale_color_manual(values =  c("limegreen","#E7B800","green4"))+
  guides(color = F, size = F)+
  labs(y="Dry above-ground biomass (g)", x="Genotype")+
  theme (legend.title= element_text(size=10), legend.text= element_text(size=10), 
       axis.title.x = element_text(size=10), axis.title.y = element_text(size=10))
```
