## Supplementary material for "Reducing chlorophyll level in seed filling stages results in higher seed nitrogen without impacting canopy carbon assimilation": SI appendix: Fig4A.R

```
#Loading packages
library(ggplot2)
library(ggpubr)
library(dplyr)

#http://www.sthda.com/english/articles/24-ggpubr-publication-ready-plots/76-add-p-values-and-significance-levels-to-ggplots/
chl<-read.csv("Fig4A.csv")

#test normality
ggqqplot(chl$Total.Chlorophyll)
shapiro.test(chl$Total.Chlorophyll)

#It is not normal distribution, thus wilcoxon rank test

#http://www.sthda.com/english/articles/24-ggpubr-publication-ready-plots/76-add-p-values-and-significance-levels-to-ggplots/

group_by(chl, Treatment, Genotype) %>%
  summarise(
    count = n(),
    mean = mean(Total.Chlorophyll, na.rm = TRUE),
    sd = sd(Total.Chlorophyll, na.rm = TRUE)
  )
compare_means(Total.Chlorophyll ~ Genotype, data = chl, group.by = "Treatment")

ggboxplot(chl, x = "Treatment", y = "Total.Chlorophyll", color = "Genotype", palette = c("#E7B800","#00AFBB"), 
          ylab="Total Chlorophyll (�g/\u33A0)", xlab = "Treatment (Urea, g)", add = "jitter")
```
