## Supplementary material for "Reducing chlorophyll level in seed filling stages results in higher seed nitrogen without impacting canopy carbon assimilation": SI appendix: Fig4BC.R

```
#Loading packages
library(ggplot2)
library(ggpubr)
library(dplyr)

#Read data
biomass<-read.csv("Fig4BC.csv")

#test normality
ggqqplot(biomass$Seed_weight_g)
ggqqplot(biomass$Weight_g)
shapiro.test(biomass$Seed_weight_g)
shapiro.test(biomass$Weight_g)

#Both are not normal distribution, thus I chose to use Wilcoxon rank test.
#comparing means of seed weight
group_by(biomass,Treatment, Genotype) %>%
  summarise(
    count = n(),
    mean = mean(Seed_weight_g, na.rm = TRUE),
    sd = sd(Seed_weight_g, na.rm = TRUE)
  )

compare_means(Seed_weight_g ~ Genotype, data = biomass, group.by = "Treatment")

#conclusion: I failed to find any significant difference in seed weight.

#comparing means of above ground biomass
group_by(biomass,Treatment, Genotype) %>%
  summarise(
    count = n(),
    mean = mean(Weight_g, na.rm = TRUE),
    sd = sd(Weight_g, na.rm = TRUE)
  )

compare_means(Weight_g ~ Genotype, data = biomass, group.by = "Treatment")

#conclusion: I failed to find any significant difference in above-ground biomass.
ggboxplot(biomass, x = "Treatment", y = "Seed_weight_g", color = "Genotype", palette = c("#E7B800","#00AFBB"), 
          ylab="Seed weight (g)", xlab = "Treatment (Urea, g)", add = "jitter") 

ggboxplot(biomass, x = "Treatment", y = "Weight_g", color = "Genotype", palette = c("#E7B800","#00AFBB"), 
          ylab="Above ground biomass (g)", xlab = "Treatment (Urea, g)", add = "jitter")

#Reference
#ggplot http://www.sthda.com/english/wiki/be-awesome-in-ggplot2-a-practical-guide-to-be-highly-effective-r-software-and-data-visualization
#http://www.sthda.com/english/articles/24-ggpubr-publication-ready-plots/76-add-p-values-and-significance-levels-to-ggplots/
#https://www.rdocumentation.org/packages/ggpubr/versions/0.4.0/topics/ggboxplot
```
