## Supplementary material for "Reducing chlorophyll level in seed filling stages results in higher seed nitrogen without impacting canopy carbon assimilation": SI appendix: Fig4DF.R

```
#Loading packages
library(ggplot2)
library(ggpubr)
library(dplyr)
library(MKpower)

#http://www.sthda.com/english/articles/24-ggpubr-publication-ready-plots/76-add-p-values-and-significance-levels-to-ggplots/
CN<-read.csv("Fig4DF.csv")


#test normality
ggqqplot(CN$Nitrogen_percent)
ggqqplot(CN$CN_ratio)
shapiro.test(CN$Nitrogen_percent)
shapiro.test(CN$CN_ratio)

#Both are not normal distribution, thus wilcoxon rank test

#http://www.sthda.com/english/articles/24-ggpubr-publication-ready-plots/76-add-p-values-and-significance-levels-to-ggplots/
#Nitrogen content

group_by(CN, Treatment, Genotype) %>%
  summarise(
    count = n(),
    mean = mean(Nitrogen_percent, na.rm = TRUE),
    sd = sd(Nitrogen_percent, na.rm = TRUE)
  )
compare_means(Nitrogen_percent ~ Genotype, data = CN, group.by = "Treatment")

ggboxplot(CN, x = "Treatment", y = "Nitrogen_percent", color = "Genotype", palette = c("#E7B800","#00AFBB"), 
          ylab="Nitrogen (%, g/g)", xlab = "Treatment (Urea, g)", add = "jitter") 

##Power for Wilcoxon test
##https://cran.r-project.org/web/packages/MKpower/vignettes/MKpower.html#power-and-type-i-error-simulations
rx <- function(n) rnorm(n, mean = 3.32, sd = 0.852) 
ry <- function(n) rnorm(n, mean = 3.89, sd = 0.166) 
sim.ssize.wilcox.test(rx = rx, ry = ry, n.max = 100, iter = 1000)
sim.ssize.wilcox.test(rx = rx, ry = ry, n.min = 20, n.max = 30, step.size = 1, 
                      iter = 1000, BREAK = FALSE)

#CN ratio
group_by(CN, Treatment, Genotype) %>%
  summarise(
    count = n(),
    mean = mean(CN_ratio, na.rm = TRUE),
    sd = sd(CN_ratio, na.rm = TRUE)
  )

compare_means(CN_ratio ~ Genotype, data = CN, group.by = "Treatment")
ggboxplot(CN, x = "Treatment", y = "CN_ratio", color = "Genotype", palette = c("#E7B800","#00AFBB"), 
               ylab="C/N", xlab = "Treatment (Urea, g)", add = "jitter") 

rx <- function(n) rnorm(n, mean = 14.7, sd = 0.570) 
ry <- function(n) rnorm(n, mean = 16.0, sd = 1.30) 
sim.ssize.wilcox.test(rx = rx, ry = ry, n.max = 100, iter = 1000)
sim.ssize.wilcox.test(rx = rx, ry = ry, n.min = 10, n.max = 20, step.size = 1, 
                      iter = 1000, BREAK = FALSE)

#Carbon
ggqqplot(CN$Carbon_percent)
shapiro.test(CN$Carbon_percent)

group_by(CN, Treatment, Genotype) %>%
  summarise(
    count = n(),
    mean = mean(Carbon_percent, na.rm = TRUE),
    sd = sd(Carbon_percent, na.rm = TRUE)
  )

compare_means(Carbon_percent ~ Genotype, data = CN, group.by = "Treatment")
ggboxplot(CN, x = "Treatment", y = "Carbon_percent", color = "Genotype", palette = c("#E7B800","#00AFBB"), 
          ylab="Carbon (%, g/g)", xlab = "Treatment (Urea, g)", add = "jitter") 

rx <- function(n) rnorm(n, mean = 57.5, sd = 1.11) 
ry <- function(n) rnorm(n, mean = 57.4, sd = 1.77) 
sim.ssize.wilcox.test(rx = rx, ry = ry, n.max = 100, iter = 1000)
sim.ssize.wilcox.test(rx = rx, ry = ry, n.min = 20, n.max = 30, step.size = 1, 
                      iter = 1000, BREAK = FALSE)
```
