## Supplementary material for "Reducing chlorophyll level in seed filling stages results in higher seed nitrogen without impacting canopy carbon assimilation": SI appendix: Fig5_2.R

```
#2021 SoyFACE trial
#Loading packages
library(ggplot2)
library(ggpubr)
library(dplyr)
library(car)
library(lme4)
library(lmerTest)
library(multcomp)
library(lsmeans)
library(simr)

##########################################################################################################################
#Display data points
#http://www.sthda.com/english/articles/24-ggpubr-publication-ready-plots/76-add-p-values-and-significance-levels-to-ggplots/
#read data
CN<-read.csv("Fig5_2.csv")


################################################################################################################################
#Complete Randomized block design

################################################################################################################################
#Nitrogen_percent
#ggplot with jitter
ggboxplot(CN, x = "Treatment", y = "Nitrogen_percent", color = "Treatment", size = 1.2,
          ylab="Nitrogen (%, g/g)", add = "jitter", palette = c("#E7B800","#00AFBB"))

# 1.Data analysis. 
#Is the data skewed?
boxplot(Nitrogen_percent~ Treatment, CN)

group_by(CN, Treatment) %>%
  summarise(
    n = n(),
    mean = mean(Nitrogen_percent, na.rm = TRUE),
    sd = sd(Nitrogen_percent, na.rm = TRUE),
    se = sd / sqrt(n)
  )

#Is this data balanced?
replications(Nitrogen_percent ~ Treatment + Block, data = CN)
!is.list(replications(Nitrogen_percent ~ Treatment + Block, data = CN))
interaction.plot(CN$Block, CN$Treatment, CN$Nitrogen_percent) 


#Is the interaction between random and fixed effect?
with(CN, interaction.plot(Treatment, Block, Nitrogen_percent)) 
residualPlots(lm(Nitrogen_percent ~ Block + Treatment, CN)) 
#SPAD~block interaction, no block effect


# 2.Model fitting

CN.lmer <- lmer(Nitrogen_percent ~ Treatment + (1 | Block), CN, REML = TRUE)  
anova(CN.lmer)
summary(CN.lmer)
ranova(CN.lmer)

# 3. Model evaluation
#Residuals
plot(CN.lmer)
qqnorm(resid(CN.lmer))
qqline(resid(CN.lmer))
ggqqplot(resid(CN.lmer))


# 4. Parameter estimates
summary(CN.lmer)
confint(CN.lmer)
anova(CN.lmer, type=3)

# Are there any differences?
CN.lmer <- update(CN.lmer)
summary(CN.lmer)
anova(CN.lmer, type=3) 
anova(CN.lmer, type=3, ddf = "Kenward-Roger") 


# 5. planned comparisons and pairwise post-hoc tests

summary(glht(CN.lmer, linfct = mcp(Treatment = "Tukey")))

lsmeans(CN.lmer, pairwise ~ Treatment)
summary(glht(CN.lmer, linfct = lsm(pairwise ~ Treatment)))
confint(glht(CN.lmer, linfct = lsm(pairwise ~ Treatment)))


#6. power analysis for linear mixed model (https://www.r-bloggers.com/2017/07/power-analysis-and-sample-size-calculation-for-agriculture/)
library(simr)
summary(CN.lmer)
powerSim(CN.lmer, alpha=0.05, nsim=1000)

#calculate sample size (https://rstudio-pubs-static.s3.amazonaws.com/484106_6b51212f20164fdd88cd7cce89bdef79.html)

#Changing the number of classes
model_ext_class <- extend(CN.lmer, along="Block", n=50)
model_ext_class

#Power curve 
set.seed(0)
p_curve <- powerCurve(model_ext_class, along="Block",breaks=c(4,8,12,16,20),
                      nsim=500,alpha=.05, progress=FALSE)
plot(p_curve)


##############################################################################################################################
##CN_ratio##
#ggplot with jitter
ggboxplot(CN, x = "Treatment", y = "CN_ratio", color = "Treatment", size = 1.2,
          ylab="C/N", add = "jitter", palette = c("#E7B800","#00AFBB"))

# 1.Data analysis. 
#Is the data skewed?
boxplot(CN_ratio~ Treatment, CN)

group_by(CN, Treatment) %>%
  summarise(
    n = n(),
    mean = mean(CN_ratio, na.rm = TRUE),
    sd = sd(CN_ratio, na.rm = TRUE),
    se = sd / sqrt(n)
  )

#Is this data balanced?
replications(CN_ratio ~ Treatment + Block, data = CN)
!is.list(replications(CN_ratio ~ Treatment + Block, data = CN))
interaction.plot(CN$Block, CN$Treatment, CN$CN_ratio) 


#Is the interaction between random and fixed effect?
with(CN, interaction.plot(Treatment, Block, CN_ratio)) 
residualPlots(lm(CN_ratio ~ Block + Treatment, CN)) 
#SPAD~block interaction, no block effect


# 2.Model fitting

CN.lmer <- lmer(CN_ratio ~ Treatment + (1 | Block), CN, REML = TRUE)  
anova(CN.lmer)
summary(CN.lmer)
ranova(CN.lmer)

# 3. Model evaluation
#Residuals
plot(CN.lmer)
qqnorm(resid(CN.lmer))
qqline(resid(CN.lmer))
ggqqplot(resid(CN.lmer))


# 4. Parameter estimates
summary(CN.lmer)
confint(CN.lmer)
anova(CN.lmer, type=3)

# Are there any differences?
CN.lmer <- update(CN.lmer)
summary(CN.lmer)
anova(CN.lmer, type=3) 
anova(CN.lmer, type=3, ddf = "Kenward-Roger") 


# 5. planned comparisons and pairwise post-hoc tests

summary(glht(CN.lmer, linfct = mcp(Treatment = "Tukey")))

lsmeans(CN.lmer, pairwise ~ Treatment)
summary(glht(CN.lmer, linfct = lsm(pairwise ~ Treatment)))
confint(glht(CN.lmer, linfct = lsm(pairwise ~ Treatment)))


#6. power analysis for linear mixed model (https://www.r-bloggers.com/2017/07/power-analysis-and-sample-size-calculation-for-agriculture/)
library(simr)
summary(CN.lmer)
powerSim(CN.lmer, alpha=0.05, nsim=1000)

#calculate sample size (https://rstudio-pubs-static.s3.amazonaws.com/484106_6b51212f20164fdd88cd7cce89bdef79.html)

#Changing the number of classes
model_ext_class <- extend(CN.lmer, along="Block", n=50)
model_ext_class

#Power curve 
set.seed(0)
p_curve <- powerCurve(model_ext_class, along="Block",breaks=c(4,8,12,16,20),
                      nsim=500,alpha=.05, progress=FALSE)
plot(p_curve)


##############################################################################################################################
##Carbon_percent##
#ggplot with jitter
ggboxplot(CN, x = "Treatment", y = "Carbon_percent", color = "Treatment", size = 1.2,
          ylab="Carbon (%, g/g)", add = "jitter", palette = c("#E7B800","#00AFBB"))

# 1.Data analysis. 
#Is the data skewed?
boxplot(Carbon_percent~ Treatment, CN)

group_by(CN, Treatment) %>%
  summarise(
    n = n(),
    mean = mean(Carbon_percent, na.rm = TRUE),
    sd = sd(Carbon_percent, na.rm = TRUE),
    se = sd / sqrt(n)
  )

#Is this data balanced?
replications(Carbon_percent ~ Treatment + Block, data = CN)
!is.list(replications(Carbon_percent ~ Treatment + Block, data = CN))
interaction.plot(CN$Block, CN$Treatment, CN$Carbon_percent) 


#Is the interaction between random and fixed effect?
with(CN, interaction.plot(Treatment, Block, Carbon_percent)) 
residualPlots(lm(Carbon_percent ~ Block + Treatment, CN)) 
#SPAD~block interaction, no block effect


# 2.Model fitting

CN.lmer <- lmer(Carbon_percent ~ Treatment + (1 | Block), CN, REML = TRUE)  
anova(CN.lmer)
summary(CN.lmer)

# 3. Model evaluation
#Residuals
plot(CN.lmer)
qqnorm(resid(CN.lmer))
qqline(resid(CN.lmer))
ggqqplot(resid(CN.lmer))


# 4. Parameter estimates
summary(CN.lmer)
confint(CN.lmer)
anova(CN.lmer, type=3)

# Are there any differences?
CN.lmer <- update(CN.lmer)
summary(CN.lmer)
anova(CN.lmer, type=3) 
anova(CN.lmer, type=3, ddf = "Kenward-Roger") 


# 5. planned comparisons and pairwise post-hoc tests

summary(glht(CN.lmer, linfct = mcp(Treatment = "Tukey")))

lsmeans(CN.lmer, pairwise ~ Treatment)
summary(glht(CN.lmer, linfct = lsm(pairwise ~ Treatment)))
confint(glht(CN.lmer, linfct = lsm(pairwise ~ Treatment)))


#6. power analysis for linear mixed model (https://www.r-bloggers.com/2017/07/power-analysis-and-sample-size-calculation-for-agriculture/)
library(simr)
summary(CN.lmer)
powerSim(CN.lmer, alpha=0.05, nsim=500)

#calculate sample size (https://rstudio-pubs-static.s3.amazonaws.com/484106_6b51212f20164fdd88cd7cce89bdef79.html)

#Changing the number of classes
model_ext_class <- extend(CN.lmer, along="Block", n=50)
model_ext_class

#Power curve 
set.seed(0)
p_curve <- powerCurve(model_ext_class, along="Block",breaks=c(4,8,12,16,20),
                      nsim=500,alpha=.05, progress=FALSE)
plot(p_curve)
```
