## Supplementary material for "Reducing chlorophyll level in seed filling stages results in higher seed nitrogen without impacting canopy carbon assimilation": SI appendix: SuppFig5_1.R

```
#2019 IL field trial for nitrogen
#reference

#packages that need
library(ggplot2)
library(ggpubr)
library(dplyr)
library(car)
library(lme4)
library(lsmeans)
library(multcomp)
library(lmerTest)

#read data
nitrogen<-read.csv("IL2019_Nitrogen.csv")
#subset data
nitrogen <- subset(nitrogen, Treatment=="1")

#Randomized complete block design test. ANOVA mixed model. fixed effect of block and genotype is random effect.
ggqqplot(nitrogen$N)

#lmer
nitrogen.lmer <- lmer(N ~ Genotype + (1|Block), data=nitrogen, REML=T) 
Anova(nitrogen.lmer, type = 3, white.adjust=TRUE)
ggqqplot(resid(nitrogen.lmer))
#model evaluation looks OK

#parameter estimates
summary(nitrogen.lmer)
anova(nitrogen.lmer, type=3) 
anova(nitrogen.lmer, type=3, ddf = "Kenward-Roger") 

#post-hoc test
summary(glht(nitrogen.lmer, linfct = mcp(Genotype = "Tukey")))

lsmeans(nitrogen.lmer, pairwise ~ Genotype)
summary(glht(nitrogen.lmer, linfct = lsm(pairwise ~ Genotype)))
confint(glht(nitrogen.lmer, linfct = lsm(pairwise ~ Genotype)))

#Ncomponent before SLA
#T0: No difference
#T1: TotalN, NO3, NH4, Organic N WT=mt<mt2
#T2: TotalN, NO3, NH4, Organic N WT=mt<mt2

#Ncomponent after SLA
#T0: No difference
#T1: TotalN, NO3, NH4, Organic N WT=mt<mt2
#T2: TotalN, NO3, NH4, Organic N WT=mt<mt2
```
