## Supplementary material for "Reducing chlorophyll level in seed filling stages results in higher seed nitrogen without impacting canopy carbon assimilation": SI appendix: SuppFig5_2.R

```
#2019 IL field trial for nitrogen
#reference

#packages that need
library(ggplot2)
library(ggpubr)
library(dplyr)
library(car)

#read data
nitrogen<-read.csv("IL2019_Nitrogen.csv")

#N
ggboxplot(nitrogen, x = "Treatment", y = "N", color = "Genotype", size = 1.2,
          ylab="Nitrogen (%,g/g)", xlab = "Treatment", add = "jitter", 
          palette = c("limegreen","#E7B800","green4"))

#Randomized complete block design test. ANOVA mixed model. fixed effect of block and genotype is random effect.
ggqqplot(nitrogen$N)

fit.lm = lm(N ~ Genotype + Treatment + Block, data=nitrogen)
anova(fit.lm)

#interaction between genotype and treatment
res.aov3 <- aov(N ~ Genotype*Treatment + Block, data=nitrogen)
summary(res.aov3)

#Compute two-way ANOVA test in R for unbalanced designs
my_anova <- aov(N ~ Genotype*Treatment + Block, data=nitrogen)
Anova(my_anova, type = "III")
#Genotype and Geno-trt interactions are significant

#C
ggboxplot(nitrogen, x = "Treatment", y = "C", color = "Genotype", size = 1.2,
          ylab="Carbon (%, g/g)", xlab = "Treatment", add = "jitter", 
          palette = c("limegreen","#E7B800","green4"))

#Randomized complete block design test. ANOVA mixed model. fixed effect of block and genotype is random effect.
ggqqplot(nitrogen$C)
fit.lm = lm(C ~ Genotype + Treatment + Block, data=nitrogen)
anova(fit.lm)

#interaction between genotype and treatment
res.aov3 <- aov(C ~ Genotype*Treatment + Block, data=nitrogen)
summary(res.aov3)

#Compute two-way ANOVA test in R for unbalanced designs
my_anova <- aov(C ~ Genotype*Treatment + Block, data=nitrogen)
Anova(my_anova, type = "III")

#CN.ratio
ggboxplot(nitrogen, x = "Treatment", y = "CN.ratio", color = "Genotype", size = 1.2,
          ylab="C/N", xlab = "Treatment", add = "jitter", 
          palette = c("limegreen","#E7B800","green4"))

#Randomized complete block design test. ANOVA mixed model. fixed effect of block and genotype is random effect.
ggqqplot(nitrogen$CN.ratio)
fit.lm = lm(CN.ratio ~ Genotype + Treatment + Block, data=nitrogen)
anova(fit.lm)

#interaction between genotype and treatment
res.aov3 <- aov(CN.ratio ~ Genotype*Treatment + Block, data=nitrogen)
summary(res.aov3)


#Compute two-way ANOVA test in R for unbalanced designs
my_anova <- aov(CN.ratio ~ Genotype*Treatment + Block, data=nitrogen)
Anova(my_anova, type = "III")

#OrganicN
ggboxplot(nitrogen, x = "Treatment", y = "OrganicN", color = "Genotype", size = 1.2,
          ylab="Organic N (g/m2)", xlab = "Treatment", add = "jitter", 
          palette = c("limegreen","#E7B800","green4"))

#Randomized complete block design test. ANOVA mixed model. fixed effect of block and genotype is random effect.
ggqqplot(nitrogen$OrganicN)
fit.lm = lm(OrganicN ~ Genotype + Treatment + Block, data=nitrogen)
anova(fit.lm)

#interaction between genotype and treatment
res.aov3 <- aov(OrganicN ~ Genotype*Treatment + Block, data=nitrogen)
summary(res.aov3)


#Compute two-way ANOVA test in R for unbalanced designs
my_anova <- aov(OrganicN ~ Genotype*Treatment + Block, data=nitrogen)
Anova(my_anova, type = "III")
```
