## Supplementary material for "Reducing chlorophyll level in seed filling stages results in higher seed nitrogen without impacting canopy carbon assimilation": SI appendix: SuppFig5_3.R

```
#2019 IL field trial for other traits
#reference

#packages that need
library(ggplot2)
library(ggpubr)
library(dplyr)
library(car)

#read data
other<-read.csv("IL2020_LAI.csv")

#LAI
ggboxplot(other, x = "Genotype", y = "LAI", color = "Genotype", size = 1.2,
          ylab="LAI", xlab = "Treatment", add = "jitter", 
          palette = c("limegreen","#E7B800","green4"), ylim = c(0, 6))

#Randomized complete block design test. ANOVA mixed model. fixed effect of block and genotype is random effect.
ggqqplot(other$LAI)

fit.lm = lm(LAI ~ Genotype + Treatment + Block, data=other)
anova(fit.lm)

#interaction between genotype and treatment
res.aov3 <- aov(LAI ~ Genotype.Treatment + Block, data=other)
summary(res.aov3)


#Compute two-way ANOVA test in R for unbalanced designs
my_anova <- aov(LAI ~ Genotype.Treatment + Block, data=other)
Anova(my_anova, type = "III")
#I faled to find significant difference in LAI between subjects.

#SLA
ggboxplot(other, x = "Treatment", y = "SLA", color = "Genotype", size = 1.2,
          ylab="SLA (m2/kg)", xlab = "Treatment", add = "jitter", 
          palette = c("limegreen","#E7B800","green4"),ylim = c(0, 50))

#Randomized complete block design test. ANOVA mixed model. fixed effect of block and genotype is random effect.
ggqqplot(other$SLA)

fit.lm = lm(SLA ~ Genotype.Treatment+Block, data=other)
anova(fit.lm)
ls_genotype_treatment<-lsmeans(fit.lm, ~Genotype.Treatment)

#interaction between genotype and treatment
res.aov3 <- aov(SLA ~ Genotype.Treatment + Block, data=other)
summary(res.aov3)


#Compute two-way ANOVA test in R for unbalanced designs
my_anova <- aov(SLA ~ Genotype.Treatment + Block, data=other)
Anova(my_anova, type = "III")
#There is a block effect, but I failed to find significant difference in SLA between subjects.

#compare models
library(lme4)
library(lsmeans)
data.sla.lmer <- lmer(SLA ~ Genotype.Treatment + (1|Block), data=other) 

Anova(data.sla.lmer, type = 3, white.adjust=TRUE)

#model evaluation looks OK
ggqqplot(resid(data.sla.lmer))

#parameter estimates
summary(data.sla.lmer)
confint(data.sla.lmer)
anova(data.sla.lmer)

mean_plot <-group_by(other, Plot) %>%
  summarise(
    count = n(),
    mean = mean(SLA, na.rm = TRUE),
    sd = sd(SLA, na.rm = TRUE)
  )
print(mean_plot)
compare_means(SLA ~ Plot, data = other)
```
