## Supplementary material for "Reducing chlorophyll level in seed filling stages results in higher seed nitrogen without impacting canopy carbon assimilation": SI appendix: Table1_IL2019.R

```
#Complete Randomized block design
#Loading packages
library(ggplot2)
library(ggpubr)
library(dplyr)
library(car)
library(lme4)
library(lmerTest)
library(multcomp)
library(lsmeans)

#Read data
#read data

data.rcb1 <-read.csv("IL2019_gasexchange_T2.csv", header = T, sep = ",", strip.white = T)
head(data.rcb1)
data.rcb1$Block <- as.factor(data.rcb1$Block)

# 1.Data analysis. 
#Is the data skewed?
boxplot(Aprime ~ Genotype, data.rcb1) 
#SPAD: no skew
#phiCO2: skewed

group_by(data.rcb1, Genotype) %>%
  summarise(
    n = n(),
    mean = mean(Aprime, na.rm = TRUE),
    sd = sd(Aprime, na.rm = TRUE),
    se = sd / sqrt(n)
  )

#Is this data balanced?
replications(Aprime ~ Genotype + Block, data = data.rcb1)
!is.list(replications(Aprime ~ Genotype + Block, data = data.rcb1))
interaction.plot(data.rcb1$Block, data.rcb1$Genotype, data.rcb1$Aprime) 
#SPAD: balanced
#phiCO2: balanced

#Is the interaction between random and fixed effect?
with(data.rcb1, interaction.plot(Genotype, Block, Aprime)) 
residualPlots(lm(Aprime ~ Block + Genotype, data.rcb1)) 
#SPAD~block no interaction, but block effect?
#phiCO2~block no interaction no block effect

# 2.Model fitting

data.rcb1.lmer <- lmer(Aprime ~ Genotype + (1 | Block), data.rcb1, REML = TRUE)  
data.rcb1.lmer1 <- lmer(Aprime ~ Genotype + (Genotype | Block), data.rcb1, REML = TRUE,
                        control = lmerControl(check.nobs.vs.nRE = "ignore"))  
anova(data.rcb1.lmer, data.rcb1.lmer1)


# 3. Model evaluation
#Residuals
plot(data.rcb1.lmer)
qqnorm(resid(data.rcb1.lmer))
qqline(resid(data.rcb1.lmer))
ggqqplot(resid(data.rcb1.lmer))


# 4. Parameter estimates
summary(data.rcb1.lmer)
confint(data.rcb1.lmer)
anova(data.rcb1.lmer, type=3)

# Are there any differences?
data.rcb1.lmer <- update(data.rcb1.lmer)
summary(data.rcb1.lmer)
anova(data.rcb1.lmer, type=3) 
anova(data.rcb1.lmer, type=3, ddf = "Kenward-Roger") 

# 5. planned comparisons and pairwise post-hoc tests

summary(glht(data.rcb1.lmer, linfct = mcp(Genotype = "Tukey")))

lsmeans(data.rcb1.lmer, pairwise ~ Genotype)
summary(glht(data.rcb1.lmer, linfct = lsm(pairwise ~ Genotype)))
confint(glht(data.rcb1.lmer, linfct = lsm(pairwise ~ Genotype)))
```
